# Three catalase-peroxidases promote extended stationary phase survival of *Vibrio natriegens* during ecologically relevant exposures to exogenous hydrogen peroxide

**DOI:** 10.64898/2026.09.14.751516

**Authors:** Liz D. Glasgo, Emily E. Chase, Spiridon E. Papoulis, Kennedi M. Hambrick, Luke T. Qualey, Slaybrina D.R. Raphael, Michael A. Gilchrist, Steven W. Wilhelm, Erik R. Zinser

## Abstract

*Vibrio natriegens* is an emerging model organism in laboratory and biotechnology research that is known for its fast growth rate and diverse metabolic capabilities. However, little is known about its response to oxidative stress. Reactive oxygen species (ROS) are ubiquitous stressors for most aerobic life and without mitigation can lead to cellular damage and sometimes death. *V. natriegens* is unusual amongst the *Vibrio* genus and Proteobacterial phylum in having three copies of *katG* encoding the bifunctional ROS defense enzyme catalase-peroxidase. The biological significance of having three copies of this protective gene was investigated with phylogenetics and gene inactivation studies. We determined that two of the *katG* copies arose recently via duplication and subsequent divergence. Analysis of single, double, and triple Δ*katG* constructs revealed each *katG* gene product contributes to survival during exposure to ecologically relevant concentrations of exogenous hydrogen peroxide (HOOH) under stationary phase conditions but found the genes were dispensable for HOOH resistance during exponential growth. We also demonstrated the involvement of the RpoS regulon in *V. natriegens* oxidative stress response in stationary phase. As an outcome of this investigation, we identified and repaired several spontaneous loss-of-function mutations of *rpoS* in laboratory cultures of the *V. natriegens* type strain ATCC 14048. Together, these results provide physiological and evolutionary insights into *V. natriegens* ROS response.

## INTRODUCTION

Following the Great Oxygenation Event 2.4 billion years ago, cellular life became threatened by harmful reactive oxygen species (ROS). ROS molecules [*e.g.* superoxide (O_2_^-^), hydrogen peroxide (HOOH), and hydroxyl radicals (OH•)], induce damage through oxidization and destabilization of cellular components including DNA, proteins, and lipids (1, 2). In aerobic environments, production of ROS occurs ubiquitously through biotic and abiotic processes (3). Consequently, adaptations to combat oxidative stress are diverse and widespread.

Bacterial genomes frequently encode multiple antioxidant genes with specialized function and regulation (4). Small molecules, metals, and pigments are thought to be early evolutionary oxidant defenses (2, 5–7), while stronger defenses arose that directly degrade ROS. Superoxide dismutase (SOD) converts O_2_^-^ into O_2_ and HOOH, and catalases and peroxidases neutralize HOOH (4, 8, 9). Catalases have been further grouped into two main families: HPII (*e.g.,* KatE) –monofunctional catalase enzymes, and HPI (*e.g.,* KatG) –bifunctional catalase-peroxidase enzymes. Monofunctional catalases are widely distributed across the tree of life and phylogenetic analysis has defined three distinct subgroupings. Despite structural differences, dismutation of HOOH is the predominant enzymatic activity exhibited by monofunctional catalases (10). Bifunctional catalase-peroxidases are found in bacteria, archaea, and fungi. Interestingly bifunctional catalase-peroxidases share great sequence similarity with plant peroxidases and little similarity with monofunctional catalases. They differ biochemically from monofunctional catalases by also exhibiting peroxidase activity, however the significance of the peroxidatic activity *in vivo* is unclear (11).

Responses to oxidative stress have been characterized for several bacterial model systems (12–14). The ability of *Vibrio* species to tolerate oxidative stress has been implicated as an important mechanism for survival of pathogenic species such as *V. cholerae, V. parahaemolyticus, V. vulnificus,* and *V. harveyi* (15–18). More recently, studies of *V. parahaemolyticus* identified the presence of two *katE* and two *katG* genes (19, 20). *katE*1 and *katE*2 were reported to respond to extracellular HOOH during exponential and early stationary phase (19–21), while *katG*1 and *katG*2 were found to be regulated by RpoS, and expressed during stationary phase (19, 20, 22). Additionally, the expression of each set of duplicate enzymes was found to differ, demonstrating non-redundancy of function.

*Vibrio natriegens* is a close relative of *V. parahaemolyticus*, yet few studies have reported on its response to oxidative stress, noting sensitivity to millimolar concentrations of HOOH (23, 24). Here, we report *V. natriegens* harbors three copies of the bifunctional catalase-peroxidase encoding gene, *katG,* we designate *katG1, katG2,* and *katG3*. We investigated the phylogeny and contribution of each gene product to ecologically-relevant HOOH exposures (nano- to micro-molar range). Our analysis suggested an early ancestor possessed two copies of *katG* but a more recent duplication event led to the occurrence of three *katG* copies in *Vibrio natriegens*, a rare feature within the *Vibrio* genus. We found that the evolutionary expansion to three *katG* genes plays a role during prolonged stationary phase conditions, as all three copies contribute to survival. Finally, we report that our strain and other commonly used strains of *V. natriegens* contain one of several mutations in *rpoS*, encoding the stationary phase sigma factor σ^s^ (RpoS). Construction of a “wild type” *rpoS* allele maximized oxidative stress response and revealed RpoS may regulate *katG1* and *katG2* but not *katG3*. This work provides novel insights into the function and evolution of *katG* genes in *V. natriegens* and importantly, uncovers physiologically important characteristics of this emerging model system.

## MATERIALS AND METHODS

### Bacterial strains and culturing conditions

*Vibrio natriegens* ATCC 14048 carrying plasmid pMMB*tfox* (EZ260) was used as the parent strain to generate the strains listed in Table S1. All strains were frozen and stored at -80° C in LB+10% glycerol. Cultures were routinely grown in Minimal Marine Heterotroph Medium (MHM) (25) with carbon sources (acetate or glucose) as indicated. Experiments were performed in acid-washed glassware unless otherwise stated. Glassware was soaked in 1 N HCL for at least 24 hours, then rinsed 6 times with MilliQ water before being filled with MilliQ and autoclaved. All cultures were incubated at 30° C in an orbital shaker (200-250 rpm) in the dark. Media was supplemented with 150 μg/mL ampicillin (amp), 100 μg/mL erythromycin (erm), and 250 μg/mL spectinomycin (spec), as needed.

### Mutant design and Multiplex Genome Editing by Natural Transformation (MuGENT)

Mutant constructs were created with splicing by overlap extension (SOE) PCR as previously described (26). Deletion constructs were generated with two-piece SOE and selectable markers and complementation constructs were generated with three-piece SOE. Briefly, 3 kb upstream and 3 kb downstream regions flanking the gene of interest or site of insertion were amplified. For two-piece SOE, primers added 20-23 bp homologous nucleotide sequences to the 3′ end of the upstream region and 5′ end of the downstream region. Three-piece SOE was performed similarly but included amplification of the desired insert (middle piece) with primers that added homologous nucleotide sequences to each of its ends. Point mutations in *rpoS* were targeted with three-piece SOE with slight modification. Primers did not contain an additional 20-23 bp homologous nucleotide sequence. The desired point mutations were embedded into middle piece primer sequences. The upstream reverse primer and downstream forward primer were completely homologous to the middle piece primers. Prior to transformation the resulting nucleotide sequence was confirmed by Sanger sequencing using nested primers internal to the SOE product. Complementation constructs were also constructed with three-piece SOE and replaced the *dns* 5′ UTR and open reading frame with that of the gene of interest, as previously described (25). All primers are listed in Table S2 and all PCRs were run with Phusion Plus Polymerase (Thermo Scientific) according to manufacturer’s protocols. Products were visualized on 1.5% agarose gels using Midori Green (Bulldog Bio). Amplicons were purified with QIAquick PCR purification Kit (Qiagen) and added together in and equal concentration, 50 ng each, as the template DNA for SOE PCR. SOE PCR was run as described (25). SOE products were gel purified with QIAquick Gel Extraction Kit (Qiagen) and used in transformations.

*Vibrio natriegens* pMMB*tfox* was used as the parent strain for Multiplex Genome Editing by Natural Transformation (MuGENT) and transformations were performed as previously described (26). Cells were cotransformed with two PCR products, a non-selectable product that creates a mutation of interest and an antibiotic resistance gene to maintain selection of transformants. The selectable markers conferred erm^R^ or spec^R^ and inserted into a prophage region on Chromosome 1 (25). Strains containing multiple edits were constructed through iterative rounds of MuGENT that exchanged the antibiotic markers to maintain selection of transformants. Δ*katG* mutants were selected on plates containing appropriate antibiotic and 1 mM sodium pyruvate (Sigma Aldrich) to scavenge any hydrogen peroxide in the agar (27, 28) that may hinder growth of the mutants. Colonies isolated from natural transformation were stored at –80° C in 96-well microtiter plates (25). Incorporation of the non-selective tDNA (transforming DNA) was verified by additional PCRs targeting amplification of the mutation site. Notably, incorporation of the *rpoS* SOE construct generates single nucleotide changes that would not be detectable by PCR amplicon size. Instead transformants were screened by bubbling assay (see below) and/or Sanger sequencing. Sanger sequencing was performed by the University of Tennessee Genomics Core or Eurofins Genomics.

### Cell viability and hydrogen peroxide (HOOH) assays

Parental and mutant strains of *V. natriegens* were assayed for cell viability and ability to degrade hydrogen peroxide (HOOH). Strains were inoculated from frozen stocks and incubated overnight in MHM with 1% carbon (acetate or glucose, as indicated). Cultures were diluted 100-fold and grown again for 20-22 hours. 2 mL of overnight cells were pelleted and resuspended three times in fresh medium matching experimental conditions before being inoculated into 5 mL cultures. Importantly, ampicillin was not added to HOOH degradation assays as it has been found to quench chemiluminescent signal (29), which was corroborated here as seen in Fig. S1. HOOH measurements were performed on the Orion-L Luminometer with an acridinium ester chemiluminescent method (30). Cell abundances were monitored by viable count assay on YTSS plates (per L: 5 g tryptone, 2.5 g yeast extract, 15 g Sigma Aldrich sea salts, 12 g agar), supplemented with 1 mM sodium pyruvate.

To screen for catalase activity *via* bubbling (O_2_ production), frozen stocks were streaked onto YTSS agar plates and incubated at 30° C overnight. Single colonies were picked with sterile toothpicks and transferred to a new YTSS agar plate. Plates were incubated for 48 hours at 30° C, then 30 % HOOH (Fisher Scientific) was spotted onto the bacterial colonies. Notably, the incubation time of 48 hours was critical to observe bubbling.

### Enumeration of KatG and KatE homologs

RefSeq proteomes were downloaded (File S2; remote_assemblies.csv) and hmmsearch, version 3.3.2, was used to search all proteomes for catalase (PFAM ID: PF00199.22) and peroxidases (PF00141.26) with gathering cutoffs. Data was filtered by phylum or genus to pull out Proteobacteria, *Vibrio spp.* and *Aliivibrio spp.* Hits (File S3, S4). The number of uniquely identified targets of KatE (PF00199.22) and KatG (PF00141.26) per organism accession were quantified. Values for bubble plots were generated by determining the number of KatE and KatG hits per organism and summing the number of organisms that fell into each numerical combination.

### Percent identity matrix

To compare the KatG sequences of *V. natriegens* ATCC 14048 a percent identity matrix of KatG1 (WP_020334780.1), KatG2 (WP_020335989.1), and KatG3 (WP_020333524.1) amino acid sequences was generated using Clustal Omega v2.1.

### Taxonomic relationships with genus *Vibrio* using *16S rRNA*

To elucidate taxonomic relationships within the genus *Vibrio* a 16*S rRNA* phylogeny was produced using 16S sequences curated by Refseq (31). Sequences were downloaded, collapsed at 100% using CD-HIT v. 4.8.1 (32), and all partial 16S sequences or sequences including unresolved nucleotides (*i.e.,* “N”) were removed giving a total of 177 sequences. Sequences were aligned in MAFFT v7.310 (33) with default parameters and 500 iterations. Alignment trimming was conducted using trimAl v1.4.rev15 with these gappyout algorithm (34), producing an alignment of 1430 bp. IQ-TREE v2.2.2.3 was used to produce a tree (1000 bootstrap replicates) using the built in model test selection (35). Based on BIC (Bayesian Information Criterion), a time dependent model with invariable sites and a gamma distribution with 4 discrete categories (*i.e.,* TIM3e+I+G4) was chosen. The resulting tree was based on an alignment of 1110 constant sites, 277 parsimoniously informative sites, and 93 singleton sites. The tree was visualized using iTol online tool with modifications (36).

### KatG based phylogeny tree

Refseq proteomes under the taxonomic ID 662 (File S5; remote_assemblies_TAXID662.csv) were downloaded and KatG sequences were identified as described above. Sequences were manually curated to remove partial sequences (number of sequences = 2189). Alignment and tree production was conducted in the same way as described above (*16S rRNA* gene tree), producing a protein alignment of 841 positions, with 177 constant sites, 576 parsimoniously informative sites, and 88 singleton sites. A JTT+I+G4 phylogenetic model was chosen (*i.e.,* protein time dependent model), and the resulting tree was again visualized using iTol with modifications (36).

### Alignment of RpoS sequences from *V. natriegens* isolates

Publicly available *Vibrio natriegens* genome assemblies were obtained from the NCBI Genome database to examine allelic variation in the *rpoS* locus relative to the laboratory reference strain *V. natriegens* ATCC 14048 (GenBank accession GCF_000818675.1). Genomes representing diverse geographic origins were included: CCUG 16371 (Sapelo Island, USA; GCF_001680045.1), WPAGA4 (Zhejiang Ocean University, China; GCF_022832815.1), PWH3a (Gulf of Mexico, USA; GCF_025757485.1), I4A (Iscuandé, Colombia; GCA_034480345.1), CL-2 (Odisha, India; GCF_036281675.1), and JSH01 (Jeju, South Korea; GCF_041475595.1). The *rpoS* coding sequences were extracted from each assembly and aligned using the Clustal Omega multiple sequence alignment tool (EMBL-EBI, version 1.2.4) with default parameters.

### Statistical analysis

One-way ANOVAs with multiple comparisons tests and Dunnett’s corrections were performed when comparing greater than two populations at once. Single deletion mutants were compared to the parental strain and double deletion mutants were compared to the triple deletion strain, unless otherwise stated.

We formally evaluated the distribution of KatE and KatG gene copy numbers using a set of models derived from the Obrechkoff bivariate Poisson distribution (OBPD), which can model negative association between counts, and the Holgate bivariate Poisson distribution (HBPD), which can model positive association (37, 38). In addition, we extended the standard OBPD and HBPD to include zero-modified mixture models with a parameter *δ*_0_ to test whether the (0,0) KatE and KatG count class deviated from its expected value given the rest of the data. Full description of the methods and results of our model analyses, including lattice diagrams, decision flowcharts, and forest plots of parameter estimates can be found in the Supplementary Material.

## RESULTS

### *V. natriegens* tolerance to hydrogen peroxide

Total coastal abundances of *Vibrio spp.* have been reported as 10^2^ –10^5^ cells/mL (39), and measurements of HOOH in coastal and brackish waters range from 0.1 nM –4.5 μM (40). We therefore investigated responses of *V. natriegens* to HOOH amendments with inocula and HOOH concentrations within these ranges. In Minimal Marine Heterotroph Medium (MHM) + 1 % glucose, exposures up to 800 nM HOOH caused no change in growth rate or yield for the *V. natriegens* type strain ATCC 14048 (Fig. 1A). However, cultures exposed to 10 μM and 200 μM experienced an initial lag or decrease in cell abundance, respectively, though they eventually recovered, and final cell abundances for all HOOH treatments matched the unamended control at ∼5 x 10^8^ CFU/mL (Fig. 1A). Growth of the cultures was concurrent with degradation of HOOH over time (Fig. 1B). Notably, growth on 1 % glucose resulted in a decrease in pH (from 7.6 to 5-5.5) but did not contribute to cell mortality over the course of the 13 hour assay (Fig. 1A). This pH drop likely occurred from fermentation of the glucose, yielding organic acids (sometimes referred to as overflow metabolism (41, 42)) that overwhelmed the medium buffer (1mM TAPS, kept low to prevent interference with our HOOH detection method).

**Fig 1.**
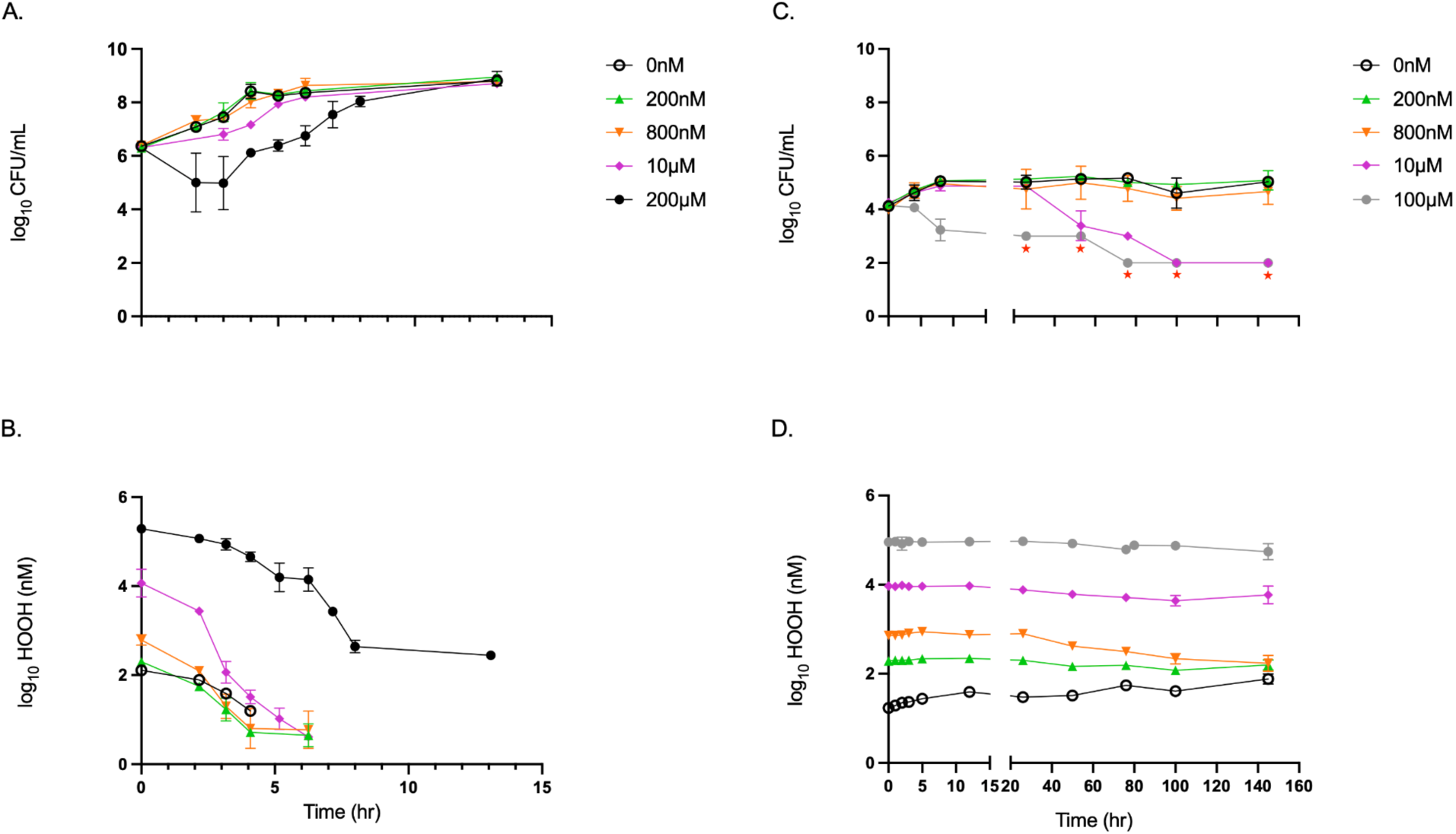
(A) Cell viability and (B) HOOH degradation of *Vibrio natriegens* ATCC 14048 in MHM + 1% glucose (1 mM TAPS). (C) Cell viability and (D) HOOH degradation of *V. natriegens* ATCC 14048 in MHM + no carbon (n=3; ± SD of the geometric mean). Red stars indicate measurements fell below the limit of detection.

The above experiments characterized the response of *V. natriegens* with replete organic carbon. To simulate resource-limited conditions that *V. natriegens* may experience in coastal and brackish waters, we repeated the HOOH exposures in MHM medium lacking additional carbon input and thus restricting growth to the trace organic contaminants of the artificial medium (43). When exposed to 0, 200, or 800 nM HOOH, final cell densities reached ∼10^5^ CFU/mL (Fig. 1C). Notably, degradation of 800 nM HOOH began after ∼24 hours and continued until ∼145 hours suggesting induction of one or more HOOH degrading enzymes after prolonged incubation (Fig. 1D). Exposure to 10 or 100 μM HOOH resulted in a loss of cell abundance below the limit of detection (<100 cells/mL) (Fig. 1C). This corresponded with little to no measurable degradation of HOOH (Fig. 1D). Importantly, limiting carbon availability established conditions where *V. natriegens* cells were resistant to ecologically relevant levels of HOOH (0 –800 nM) but became sensitive at relatively high concentrations (10 –100 μM). These results suggest growth substrate and/or cellular abundance influence the response of *V. natriegens* to HOOH and help to constrain conditions which detect a sensitivity to HOOH.

### Richness of catalase enzymes in *V. natriegens*

The robust response of *V. natriegens* to hydrogen peroxide prompted us to explore the genetic underpinnings of its oxidative stress response, beginning with catalase (KatE) and catalase-peroxidase (KatG). Diversity and richness of oxidative stress response enzymes can vary widely across bacterial phyla (8, 44). Strikingly, *V. natriegens* ATCC 14048 lacks a *katE* homolog but has three copies of *katG* located on Chromosome 2 of *V. natriegens* ATCC 14048. While other *Vibrio spp.* have been reported to carry up to two copies of *katG* or *katE* (20, 22), to our knowledge this is the first report of a *Vibrio* species with more than two copies of *katG*.

We designated the three *katG* genes of *V. natriegens “katG1”, “katG2”, and “katG3”* based on their relative positions from the origin on reference genome (NCBI Refseq accession NZ_CP016346: locus tags BA890_RS15465, BA890_RS16825, and BA890_RS19750, respectively). Protein sequence alignments of KatG1 (WP_020334780.1), KatG2 (WP_020335989.1), and KatG3 (WP_020333524.1) revealed that KatG2 and KatG3 share the highest amino acid percent identity at 88.93%, while KatG1 shares 80.36% and 78.67% amino acid identity with KatG2 and KatG3, respectively (Table 1). These results indicate a particularly high divergence of KatG1 with KatG2 and KatG3.

**Table 1.** Clustal Omega alignment percent identity matrix of *Vibrio natriegens* ATCC 14048 KatG amino acid sequences. KatG1 = WP_020334780.1 (723 amino acids), KatG2 = WP_020335989.1 (724 amino acids), KatG3 = WP_020333524.1 (723 amino acids).

|  | KatG1 | KatG2 | KatG3 |
| --- | --- | --- | --- |
| KatG1 | 100 | 80.4 | 78.7 |
| KatG2 | 80.4 | 100 | 88.9 |
| KatG3 | 78.7 | 88.9 | 100 |

The evolutionary history of the three KatG copies of *V. natriegens* was explored by constructing phylogenetic trees for the entire *Vibrio* genus (Fig. 3, S2). Sequences from a manually curated set of KatG proteins were collapsed at 100% and each branch represents a unique KatG sequence. Twelve species were identified with more than one phylogenetically distinct KatG sequence. The divergence of KatG sequences within a single species can be observed by the distance between individual or clusters of KatG sequences from the same species, denoted by color strips (Fig. 3, S2A). As a general trend, KatG sequences from different species are more closely related than KatG’s within the same species or even closely related species, the latter indicated by comparison to the *16S rRNA* gene tree (Fig. S2B). However, the sequence similarity and close phylogenetic distance between KatG2 and KatG3 in *V. natriegens* suggests recent duplication and divergence events were responsible for the establishment of the three KatG copies in this species (Fig. 3).

We next addressed whether *V. natriegens* is unusual within the *Vibrio* (and *Allivibrio*) genus and within all Proteobacteria for carrying three copies of KatG. The number of KatG (PF00141.26) and KatE (PF00199.22) homologs in all Proteobacteria and within *Vibrio* and *Allivibrio spp*. were quantified using available RefSeq proteomic sequence data. Analysis of all Proteobacterial species revealed a wide distribution in the number of KatE and KatG homologs per proteome, reaching up to nine KatE copies and five KatG copies in a single species (Fig. S3). However, most organisms contained only 1 –2 copies of each enzyme. Similar trends were observed at the genus level for *Vibrio* (and *Allivibrio*): the number of KatE and KatG homologs per proteome ranged between 1 –3, with most species containing a single copy of one or both enzymes (Fig. 2). Besides *V. natriegens, V. nitrifigilis* was the only other *Vibrio* species containing three copies of KatG and zero copies of KatE.

**Fig 2.**
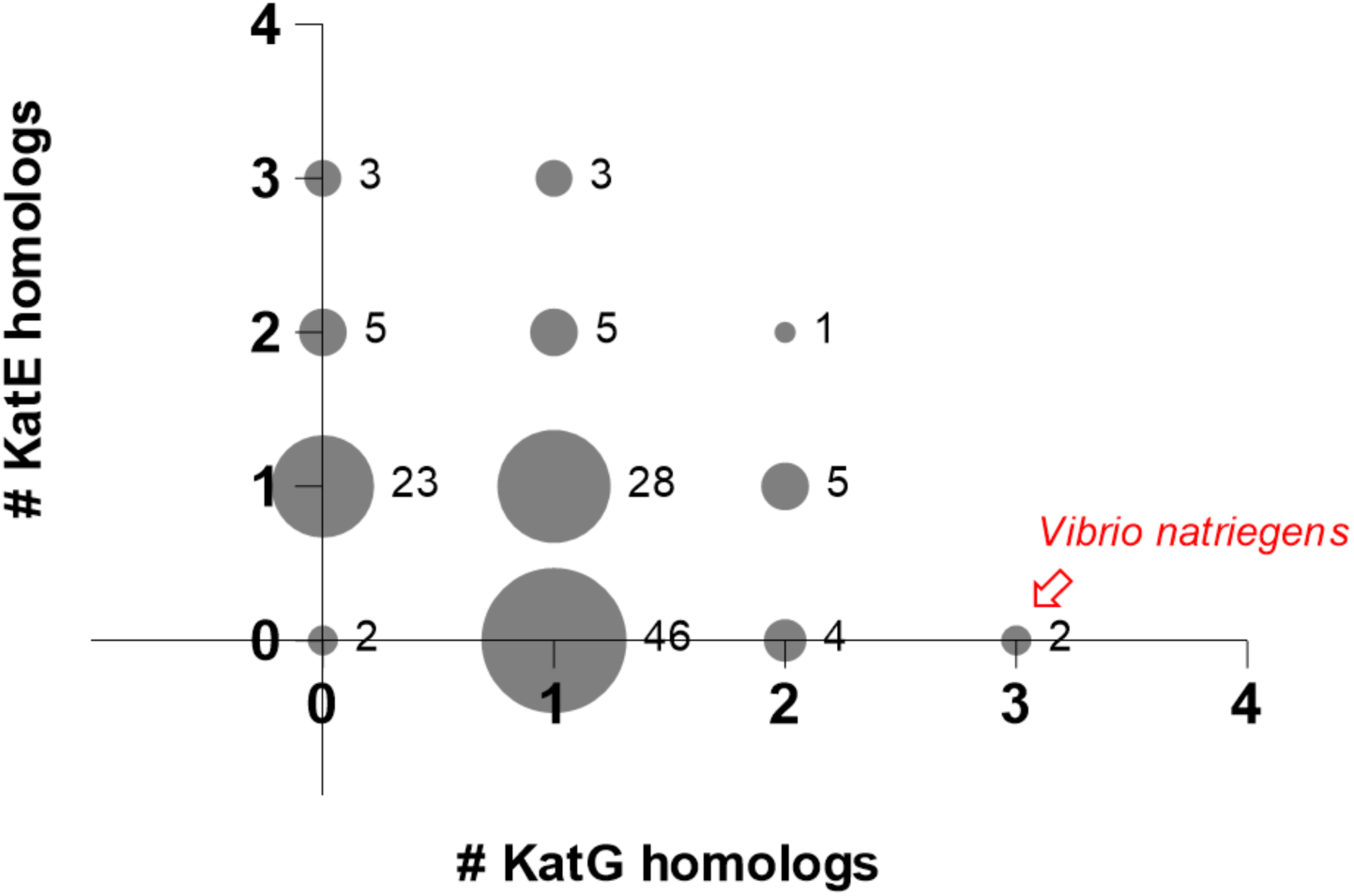
Enumeration of KatE (PF00199.22) and KatG (PF00141.26) within *Vibrio spp.* and *Aliivibrio spp.* proteomes (Refseq proteomes accessed February 2022).

**Fig 3.**
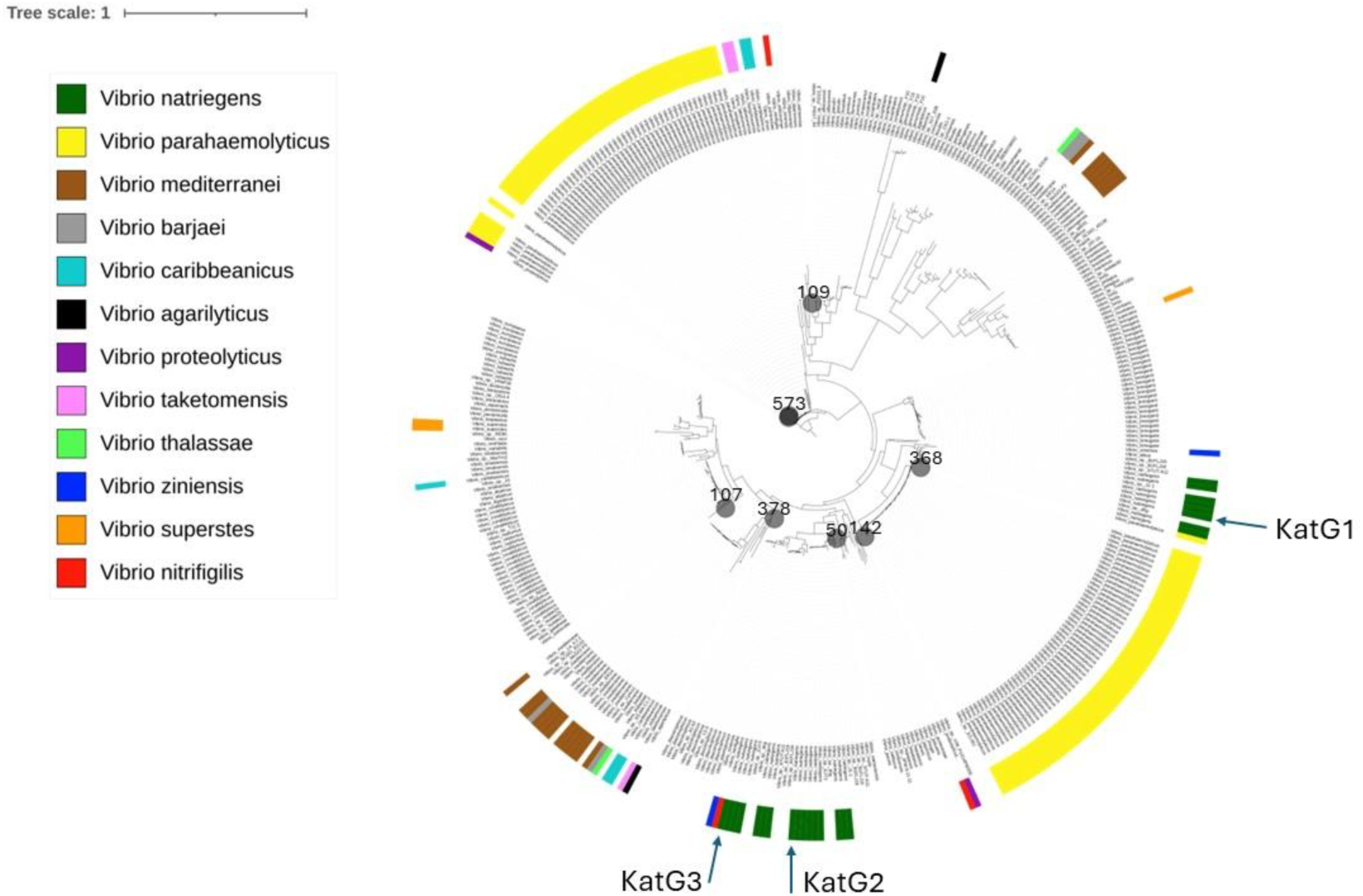
*Vibrio spp.* KatG based phylogeny. Species containing greater than one KatG (PF00141.26) copies are indicated by color strips. *V. natriegens* ATCC 14048 KatG proteins indicated by blue arrows (KatG1 = WP_020334780.1, KatG2 = WP_020335989.1, KatG3 = WP_020333524.1.). Collapsed branches are represented by grey circles and the number of collapsed branches within each circle is indicated.

KatE is common amongst the Proteobacteria and *Vibrio* genus specifically, yet *V. natriegens* lacks this enzyme. At first glance, the data from Figure S1 and Figure 2 might suggest a negative relationship between the number of copies of KatE and KatG, perhaps revealing an important interaction between the two classes of enzyme and helping to explain why *V. natriegens* lacks KatE. To address this possibility, we formally evaluated the distribution of KatE and KatG copy numbers using a set of models derived from the Obrechkoff bivariate Poisson distribution (OBPD), which can model negative association between counts (37), and the Holgate bivariate Poisson distribution (HBPD), which can model positive association (38). After accounting for the significant underrepresentation of genomes lacking both KatE and KatG (see Methods and Supplemental Material), we found no evidence that KatE and KatG counts are positively or negatively associated under either model (Obrechkoff: 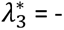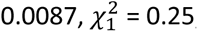, p = 0.62; Holgate: λ_3_ = 0.0000, 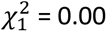, p = 1.00). The best-supported OBPD model (Table S3) has KatE and KatG counts statistically independent 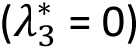.

### Impact of growth phase on Δ*katG* mutant responses to HOOH

While informatics indicated *V. natriegens* has an unusually high number of KatG copies, whether each copy contributes to fitness had yet to be addressed. Therefore, single, double, and triple Δ*katG* mutants were constructed to assess the protective role of each *katG* gene product under different nutrient conditions and exposures to HOOH. Importantly, in the absence of exogenous HOOH, no mutant strains exhibited defects in growth rate or yield (Fig. S4).

The protective role of each *katG* gene of *V. natriegens* was assessed during lag, exponential growth, and early stationary phase in MHM + 24 μM acetate medium. This medium enabled growth via aerobic respiration to final yield of ∼10^6^ cells/mL. Parental and Δ*katG* strains were first grown for 24 hours in MHM + 1 % acetate, then washed and inoculated at ∼10^3^ cells/mL in MHM + 24 μM acetate with or without HOOH amendment. Mutants lacking one, two, or three *katG* genes exhibited no defects in growth or HOOH degradation during lag and exponential growth phase when exposed to 1400 nM HOOH (Fig. S5). Notably, for all strains significant HOOH degradation did not occur within the first nine hours of growth. However, degradation was complete by 30 hours, after significant population growth and entry into stationary phase occurred.

To investigate the HOOH dynamics in stationary phase specifically, cultures were inoculated in MHM + 24 μM acetate as before but not exposed to 1400 nM until 24 hours post inoculation. All genotypes experienced an order of magnitude drop in viable counts during the first 24 hours of stationary phase, even the parental strain without the HOOH amendment (Fig. 4A and 4C). After this initial decline, viable counts stabilized for the parental strain and all *katG* mutants including the triple deletion, showing no clear protective effect for any of the *katG* copies.

**Fig 4.**
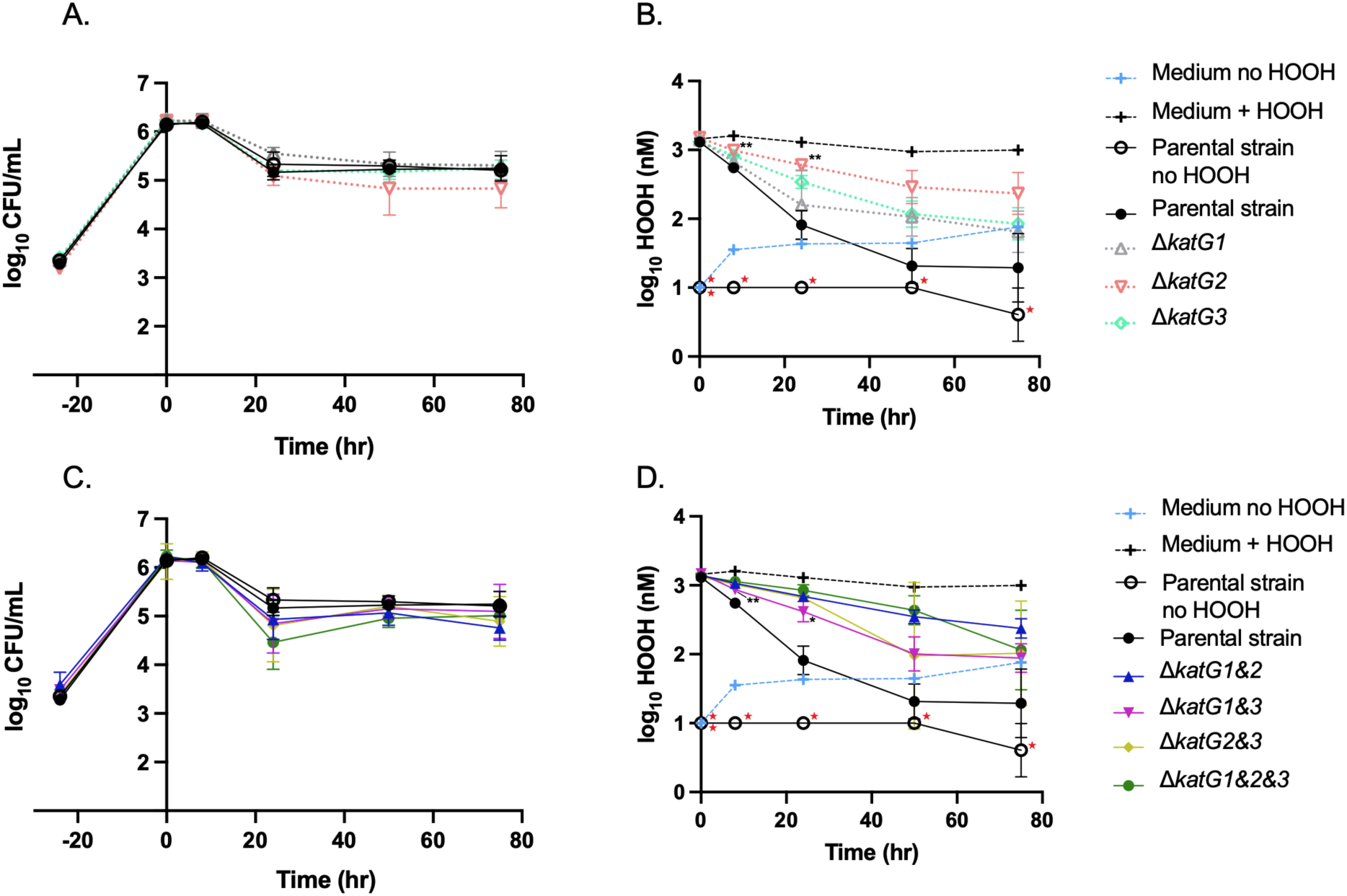
(A and C) Cell viability and (B and D) HOOH degradation by *V. natriegens* parental and mutant strains in MHM + 24 μM acetate (n=3; ± SD of the geometric mean). HOOH exposure at time 0, cells in stationary phase. P-values were calculated using One-way ANOVA multiple comparisons with Dunnet’s correction, * P ≤ 0.05, ** P ≤ 0.01: (B) Single *katG* deletions compared to parental strain, (D) double deletions compared to triple *katG* deletion. Red stars indicate measurements fell below the limit of detection. The same data for parental strain and medium +/- HOOH are shown in panels A + C and B + D to facilitate direct comparisons with the other (deletion strain) data within each panel.

However, significant differences were observed in HOOH degradation rates for some mutants during this stationary phase period (Fig. 4B and 4D). First, single knockout mutants were compared to the parental strain. The Δ*katG2* strain (EZ265) had significantly decreased ability to degrade HOOH at early time points (Fig. 4B). While not significant, this trend continued through later time points and Δ*katG2* cultures ended with higher HOOH concentrations. Likewise, differences between the parental and Δ*katG1* (EZ264) or Δ*katG3* (EZ266) strains were non-significant, however, early time points trend with higher HOOH concentrations.

To assess the contribution of each individual *katG* gene to HOOH degradation, *katG* double knockouts with only a single *katG* remaining were compared to the triple mutant lacking all *katG* copies. A trend of higher HOOH degradation was observed for all three variations of *katG* double knockouts relative to the triple knockout, suggesting each *katG* in isolation can contribute to degradation (Fig. 4D). However, only Δ*katG1&3* (EZ268) differed significantly from Δ*katG1&2&3* (EZ270), and only at early time points (Fig. 4D). This indicates that KatG2 significantly contributes to HOOH degradation under these conditions. Together, the results from all mutant genotypes suggest that all three KatG’s and especially KatG2 are active and degrade HOOH during stationary phase. Notably, the triple mutant lacking all three KatG’s maintained significant HOOH degradation activity, suggesting peroxidases are also functioning under these early stationary phase conditions.

Exposing *V. natriegens* to extended stationary phase revealed a greater dependency on the three catalase-peroxidases for survival. In this variation of the experiment, cells from the stationary phase cultures grown in MHM + 1% acetate were inoculated into MHM without a carbon source and incubated 24 hours prior to the 1400 nM HOOH addition. The parental strain was completely resistant to the HOOH exposure in extended stationary phase conditions (Fig. 5A), but left ∼400 nM HOOH behind (Fig. 5B), in contrast to its ability to more completely degrade HOOH in early stationary phase (Fig 4D). Notably, the triple *katG* mutant exhibited no ability to remove HOOH during extended stationary phase (Fig. 5D). This contrasts with its less severe phenotype in early stationary phase (Fig. 4D) and suggests that the catalase-peroxidases were the only enzymes that could scavenge exogenous HOOH during prolonged starvation (Fig. 5D).

**Fig 5.**
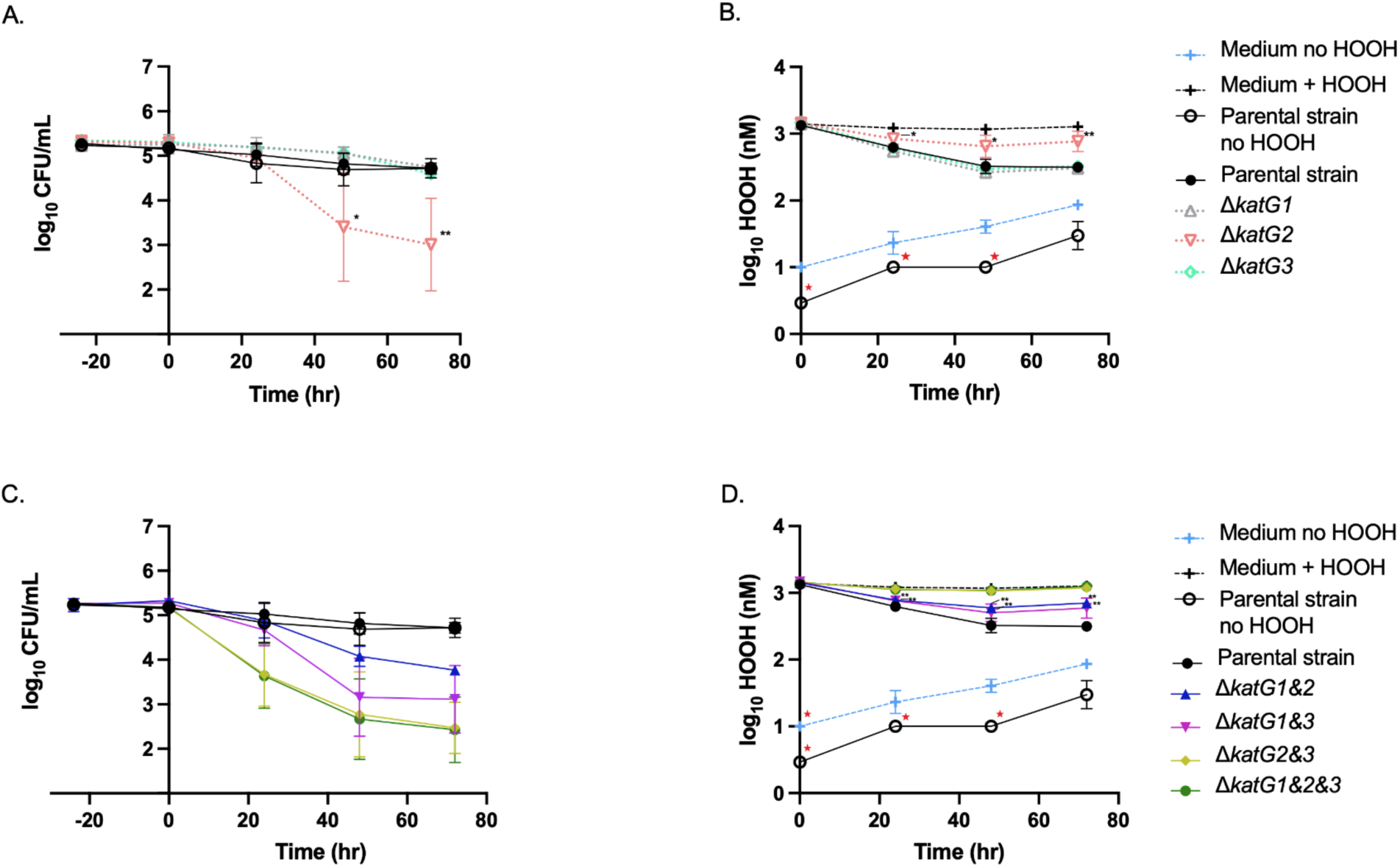
(A and C) Cell viability and (B and D) HOOH degradation by *V. natriegens* parental and mutant strains in MHM + no carbon (n=3; ± SD of the geometric mean). HOOH exposure at time 0, after 24 hours of starvation. P-values were calculated using One-way ANOVA multiple comparisons with Dunnet’s correction, * P ≤ 0.05, ** P ≤ 0.01: (B) Single *katG* deletions compared to parental strain, (D) double deletions compared to triple *katG* deletion. Red stars indicate measurements fell below the limit of detection. The same data for parental strain and medium +/- HOOH are shown in panels A + C and B + D to facilitate direct comparisons with the other (deletion strain) data within each panel.

During extended stationary phase conditions two of the *katG* genes were functional and contributed to survival of HOOH exposure. The singular knockout *ΔkatG2* exhibited slower HOOH decay and a decline in cell viability by 48 hours post exposure (Fig. 5A and 5B), which is consistent with but more pronounced than its phenotype in early stationary phase (Fig. 4A and 4B). Double knockout mutants *ΔkatG1&2* (*katG3*+) and *ΔkatG1&3* (*katG2*+) had lower HOOH decay and higher death compared to the parental strain, but significantly higher HOOH decay and less death than the triple mutant lacking all catalases. In contrast, *ΔkatG2&3* (*katG1+*) cells were indistinguishable from the triple mutant, suggesting *katG1* is not contributing to HOOH removal and cell survival under these conditions (but see section on *rpoS*, below). However, comparison of Δ*katG3* to Δ*katG1&3* suggests that the presence of *katG1* does improve cell viability and HOOH degradation when working in combination with *katG2*. Notably, these results demonstrate that during nutrient starvation any single *katG* gene is insufficient to maintain parental levels of cell viability or HOOH degradation.

### RpoS control of two *katG* genes in *Vibrio natriegens*

The sigma factor RpoS is a global regulator of stress response and stationary phase genes and is a known regulator of *katG* in several bacterial species (22, 45). To assess if RpoS likewise regulates any of the *katG* copies in *V. natriegens* we sought to genetically manipulate the *rpoS* gene. However, genome sequencing of our copy of *Vibrio natriegens* ATCC 14048 (EZ260), from which all strains in this study were derived, revealed two single nucleotide polymorphisms in *rpoS* relative to published NCBI accessions (e.g. NZ_CP160345 and NZ_CP009977): a nonsense mutation relatively early in the open reading frame (E71→ *, GAA→TAA) and a missense mutation (N123 → I, AAC → ATC). That these two polymorphisms are also present in the genome provided on the ATCC website for this strain (sequenced August 27, 2019) suggests these polymorphisms were already present when the strain was purchased.

Hypothesizing that the nonsense mutation likely rendered RpoS null in our strains we constructed various combinations of the two polymorphisms. During strain construction we noted that only the combination of E71 + I123 provided maximal “bubbling” (*i.e.,* oxygen generation via catalase activity) when HOOH was pipetted onto colonies (Video S1, Fig. S6, and see Materials and Methods). The NCBI allele E71 + N123 provided marginal bubbling and the ATCC allele of *71 + I123 provided almost no bubbling. These bubbling phenotypes correlate with the phenotypes in liquid culture (see below). Notably, six other isolates of *V. natriegens* from wide-ranging environments all carried the E71+I123 maximum bubbling allele (Fig. S7). Given that these initial observations suggest the E71+I123 combination conferring maximum activity is the prevalent allele in nature, we denote it as wild type (WT), and suggest that the two other alleles arose via lab-acquired mutations prior to sequencing for NCBI or ATCC (Fig. S8).

Next, we repeated the HOOH exposure during extended stationary phase experiments with the E71+I123 (WT) allele and E71+N123 allele of *rpoS*. Parental with WT *rpoS* cultures fully degraded the HOOH and showed no loss in viable counts (Fig. 6A and 6B), a noted improvement compared to the nonsense *rpoS* mutant (Fig. 5B). In the WT *rpoS* background, survival during HOOH exposure improved for *ΔkatG1&2&3* but still showed significant loss in viable counts by the end of incubation (Fig. 6A). Presence of any of the three *katG* genes significantly improved viability, indicating they all played a protective role in this condition (Fig. 6A).

**Fig 6.**
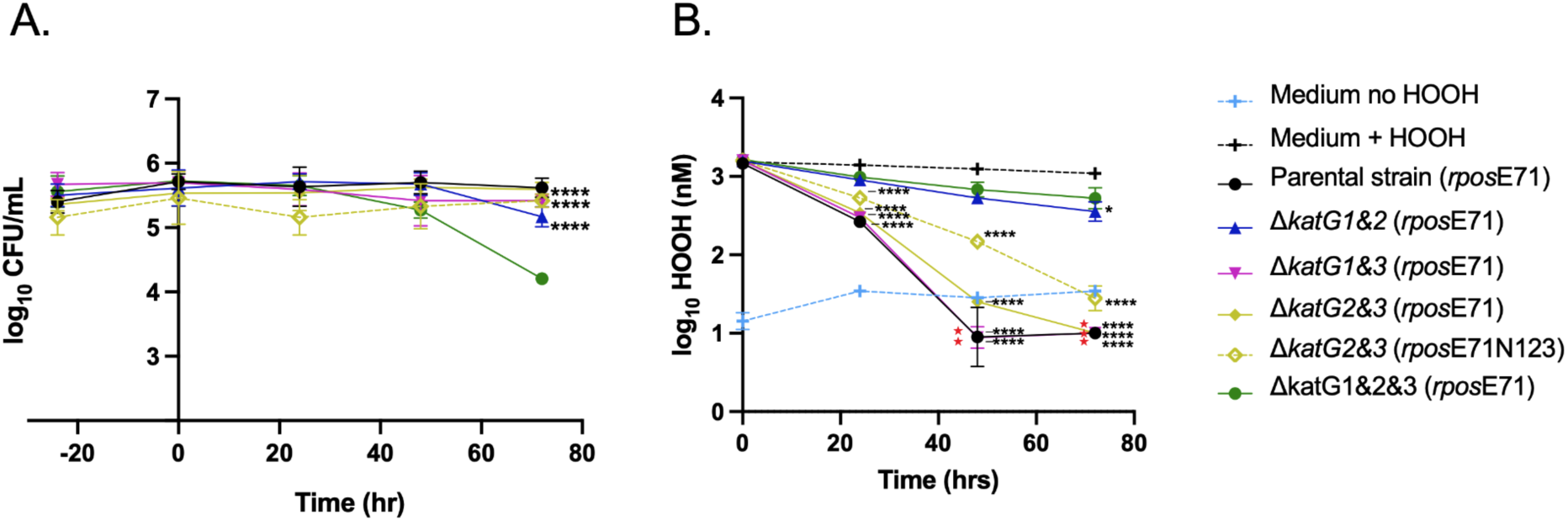
(A) Cell viability and (B) HOOH degradation by *V. natriegens* parental and mutant strains with indicated *rpoS* alleles (n=3; ± SD of the geometric mean). HOOH exposure at time 0, after 24 hours of starvation in MHM + no carbon. P-values were calculated using One-way ANOVA multiple comparisons with Dunnet’s correction, * P ≤ 0.05, ** P ≤ 0.01, **** P ≤ 0.0001: double *katG* deletions compared to triple *katG* deletion. Red stars indicate measurements fell below the limit of detectio

Importantly, *rpoS* allelic state had a significant impact on the activity of the KatG1 enzyme when working in solo. This is evident by the significant decay of HOOH in the wild type *rpoS* background (Fig. 6B) compared to non-decay in the nonsense mutant background (Fig. 5D) for the Δ*katG2&3* double mutant. Of note, Δ*katG2&3 rpoS*E71N123 demonstrated slower HOOH degradation than Δ*katG2&3 rpoS*E71I123 (WT), consistent with the bubbling assay.

HOOH degradation by Δ*katG1&3* (*katG2*+) was improved in the *rpoS* WT background and was indistinguishable from the *rpoS* WT strain containing all three *katG* genes (Fig. 6B). In the *rpoS* WT background, HOOH degradation by Δ*katG1&2* (*katG3+*) showed marginal improvement relative to the triple *katG* mutant that was less pronounced than in the *rpoS* nonsense mutant background (Fig. 5D). Finally, we noted greater degradation of HOOH by Δ*katG1&2&3* in WT *rpoS* (Fig. 6B) than in the nonsense *rpoS* background (Fig. 5D), consistent with RpoS control of HOOH-degrading enzymes other than the three KatG’s.

Phenotypic complementation of *katG1, katG2,* or *katG3* deletion was tested by placement of a gene copy with native promoter on the chromosome rather than on a multicopy plasmid; this was done to minimize artefactual elevated activity due to overexpression of any one of these single copies of *katG*. Copies were inserted ectopically into the *dns* locus of Δ*katG1&2&3 rpoS*E71I123 creating strains with single *katG* genes in one copy per chromosome; Δ*katG1&2&3 rpoS*E71I123 *Δdns:: katG1* (EZ310), Δ*katG1&2&3 rpoS*E71I123 *Δdns:: katG2* (EZ311), and Δ*katG1&2&3 rpoS*E71I123 *Δdns:: katG3* (EZ312). HOOH exposure during prolonged stationary phase was performed with the complement strains, each demonstrated successful restoration of HOOH degradation and complementation of the *katG* genes (Fig. S9).

## DISCUSSION

This study examined the evolutionary history and the ecologically relevant functions of the three catalase-peroxidases of *V. natriegens* ATCC 14048. The presence of three catalase-peroxidase genes (*katG1,2,3*) is rare amongst *Vibrio* species and likely a consequence of two independent acquisition events followed by a duplication and divergence of one of the acquired copies. All three *katG* genes are conditionally required for full protection during early and prolonged stationary phase conditions but are dispensable during exponential growth. This work also led to the identification of two different *rpoS* point mutations in frequently cited laboratory strains of ATCC 14048 and repair of the wild type *rpoS* led to improved HOOH survival during starvation, potentially through the regulation of *katG1* and *katG2*.

### Catalase-peroxidases are important during stationary phase but not log phase

Stationary phase cultures of *E. coli* and other microbes possess high catalase activity (19, 20, 46–49), and our study suggests a similar response by *V. natriegens*. We observed that all three copies of catalase-peroxidase were required for cells to survive ecologically relevant concentrations of HOOH under nutrient starvation conditions. Stationary phase is defined as a period of (imperfect) survival in absence of net population growth, and management of co-occurring stresses is a well-established paradigm. Our studies indicate that like other species, *V. natriegens* cells entering stationary phase are primed for oxidative stress resistance through expression of catalases, and this priming contributes to the survival of this microbe during periods of starvation in its coastal/estuarine habitat.

While essential for stationary phase, all three *katG* genes appear dispensable for HOOH resistance during exponential growth in liquid batch cultures. Interestingly, cell cultures exposed to HOOH during lag phase demonstrated unaltered growth and complete HOOH degradation by 30 hours. Although cell growth appeared normal during the first nine hours after inoculation of batch cultures, there was a curious absence in HOOH degradation during this time. It is possible that actively growing and dividing cells of *V. natriegens* mitigate damage through other means such as DNA repair mechanisms and/or asymmetric division of damage to daughter cells (50, 51). This would explain why growth was not impacted in absence of HOOH degradation. Alternatively, cells could have been degrading enough HOOH to keep intracellular levels low during the first nine hours but the degradation was not detectable in the medium until a higher cellular concentration was reached.

### Evolutionary acquisition of three catalase-peroxidases

While not unique amongst the *Vibrio* genus or the Proteobacteria, acquisition of three copies of *katG* (and no *katE*’s) is certainly unusual (Figures 2 and S1). Why *V. natriegens* took this evolutionary pathway is a matter of speculation. However, if we assume selection rather than drift as the evolutionary driver, we note that all three copies contribute to survival under oxidative stress conditions during prolonged stationary phase, conditions that may be common for *V. natriegens* in its aquatic habitat. Selection under these specific conditions would thus favor the acquisition of all three copies, as strains with all three copies survives better than strains with any combination of two copies.

The presence of multiple catalases per genome is common in Proteobacterial and *Vibrio* species (Fig. 2 and Fig. S1) but in most cases this involves a combination of KatE (monofunctional catalase) and KatG (bifunctional catalase-peroxidase). This raises the question of why all three *V. natriegens* catalases are KatGs and not a combination of KatG and KatE. Our formal analysis with a zero-modified extension of Obrechkoff bivariate Poisson mixture model indicated that amongst the *Vibrio* genus and the Proteobacteria as a whole, the number of KatGs per genome were independent of the number of KatEs per genome. Thus, it would appear that the existence of one or more KatGs in the *V. natriegens* genome did not preclude the acquisition of a KatE. However, we note the important caveat that our analysis ignores the shared ancestry underlying the data, and thus our results cannot be used to make inferences about evolutionary differences between these groups. A deeper analysis incorporating shared ancestry should provide a more robust understanding of the relationships between KatG and KatE distributions.

Despite the uncertainties stated above, we suspect that the dual activity of the KatG’s provide greater fitness to *V. natriegens* in its native environment relative to the monofunctional KatE’s. Catalases degrade HOOH in two steps, and the second step can stall when HOOH concentrations are low (11). For monofunctional catalases, it has been proposed that binding to NADP(H) may help to protect stalled enzymes from inactivation, but more work is necessary to support this idea (52). By comparison, peroxidase function of *katG* can complete the second step even at low concentrations of HOOH. While the decay rates from the KatG’s peroxidase function are slow relative to (monofunctional) peroxidase enzymes also present in the cell (9, 11), this catalase rescue pathway may prove important for microbes exposed to low as well as high concentrations of HOOH that would be expected for coastal and estuarine environments. Interestingly, a recent survey of catalases in marine systems suggested that KatE’s are more prevalent in bacteria that associate with particulate organic carbon versus bacteria that have more of a free-living lifestyle (53). Given the ability of *V. natriegens* to associate with solid surfaces [this study and (54)], future studies on the microscale *in situ* distributions of this bacterium should be informative with respect to its catalase composition.

### *rpoS* role in oxidative stress and its mutation in laboratory cultures

The role of RpoS in the stress response of *V. natriegens* is largely unexplored, though a recent study suggested a minor role in acid resistance (55). To our knowledge this is the first study of RpoS function in the oxidative stress response of *V. natriegens*. We observed that like the *katG*s, *rpoS* is dispensable for HOOH resistance during exponential growth in liquid batch cultures and biofilm formation but important for survival in stationary phase. Whether RpoS regulates transcription of the three *katG* copies in *V. natriegens* and if this regulation is direct or indirect (i.e., through an RpoS- controlled regulator of *katG*) requires additional experimentation beyond the scope of this study. However, we note that reconstruction of the wild type *rpoS* allele –as assessed by HOOH degradation activity –in different *katG* backgrounds improved HOOH defense and ultimately suggested *katG1* and *katG2* may be under RpoS control. Additionally, in mutants lacking all three catalase-peroxidases survival in extended stationary phase in the presence of HOOH improved dramatically (Fig. 6), suggesting that the RpoS regulon involves protection beyond the three catalase-peroxidases. As a final note, prior studies (in the RpoS nonsense background) from our group (25) and another (23) implicated the transcription regulator OxyR in protecting *V. natriegens* from oxidative stress. Thus, a more extensive and focused investigation of both OxyR and RpoS in the regulation of KatG expression and the oxidative stress response in this species is certainly warranted.

Loss of function mutations in *rpoS* may occur frequently in laboratory cultures of *V. natriegens* and this may extend beyond strain ATCC 14048. A previous study reported that *V. natriegens* strain CCUG 16347 has low intrinsic catalase activity informed by the lack of bubbling when spotted with HOOH and a cold temperature sensitivity that could be rescued by plating cells with exogenous catalase (24). An investigation of the published genome of that strain revealed that the *rpoS* gene contains a stop codon mutation at amino acid 59 due to an upstream frameshift mutation (FS S49/L50 (+C)). Notably, this is a different nonsense mutation than found in ATCC 14048, but may likewise be responsible for the low catalase activity in the strain. Additionally, loss of function mutations in *rpoS* have been reported extensively for lab cultures of other species such as *E. coli*. Storage and transport conditions are credited for unintentional selection of *rpoS* mutants in *E. coli* collections and mutations in *rpoS* can provide a fitness advantage (56). Notably, surveys of fresh environmental isolates indicate that *rpoS* mutations are rare in natural *E. coli* populations (57, 58). In *V. natriegens* the diversity of *rpoS* alleles and frequency of mutations is not well constrained, but in notable contrast to the (unrepaired) ATCC 14048 strain, 6 other *V. natriegens* isolates carried the E71 + I123 ‘wild-type’ allele (Fig. S7), suggesting loss of function mutations may likewise be rare for this species.

As a final observation, the extent of extracellular HOOH degradation exhibited condition and genotype dependence. In the *rpoS* nonsense mutant background, complete degradation occurred when cells were exposed during early stationary phase, whereas roughly 400 nM HOOH was left behind when exposed during extended stationary phase (Fig. 4 and 5, respectively). A potential explanation for the incomplete HOOH degradation is that cells in extended stationary phase lack significant reductant to operate peroxidases and thus must rely exclusively on catalase activity, which has the potential to stall at low HOOH concentrations (13). We note however that restoration of the WT *rpoS* allele facilitated complete HOOH degradation, perhaps through regulation of antioxidant enzymes and/or metabolic changes that provide reductant for the peroxidases.

## Conclusions

Our investigation of the catalase-peroxidases contributes to a new understanding of the oxidative stress response of *V. natriegens.* We have discovered that an unusually high number of *katG* copies is required for maximal protection from ecologically-relevant concentrations of exogenous HOOH under some but not all states of growth. These results have implications for both the cultivation and ecological and evolutionary understanding of this fast growing, industrially important microbe. Importantly, we highlight mutations in *rpoS* among culture stocks of *V. natriegens* that were most likely acquired in the laboratory during serial passage. This observation may prove valuable for future research in *V. natriegens* physiology and genetics.

## ACKNOWLEDGEMENTS

This work was supported by a Student/Faculty Research Award from the University of Tennessee, Knoxville to L.D. Glasgo and E.R.Z and NSF OCE-2023680 to E.R.Z. A portion of the computation for this work was performed on the University of Tennessee Infrastructure for Scientific Applications and Advanced Computing (ISAAC) computational resources. The authors would like to thank the High Performance & Scientific Computing (HPSC) group (UTK) for their support. We’d also like to thank Liz Fozo, Zachary Slimak and Joan Slonczewski for valuable discussions

