## Supplemental File 1 for "Three catalase-peroxidases promote extended stationary phase survival of *Vibrio natriegens* during ecologically relevant exposures to exogenous hydrogen peroxide"

**Supplemental Material**

**Supplemental methods and analysis:**

We wanted to quantitatively evaluate the relationship between copies of the *katE* and *katG* genes per genome observed across taxa (**Supplemental File 3 and 4** for *Vibrio* genus and Proteobacteria, respectively). Given the gene count data is by its nature non-negative and discrete combined with the relatively low gene copy number in any given genome (median = 1, max = 9), and because we are interested in determining if there are any negative or positive association between gene counts, the data is best modeled using a bivariate Poisson distribution (BPD). Unfortunately, there is no well-defined BPD that models both positive and negative associations between counts, so we utilized two different BPD models. Specifically, we used the Obrechkoff BPD (OBPD; Ghosh *et al.* 2021) to test for a negative association between KatE and KatG counts across species and the Holgate BPD (HBPD; Holgate 1964) to test for a positive association between counts. It is important to recognize that our analysis treats each set of KatE and KatG counts as an independent observation. In other words, our analysis fails to account for shared phylogenetic history, making our tests descriptive in nature rather than explicitly testing different evolutionary hypotheses about KatE and KatG evolution.

With that caveat in mind, the standard OBPD uses three parameters: $\lambda_1$ and $\lambda_2$ represent the univariate Poisson rate parameters for KatE and KatG, respectively, when their counts are independent of one another. The third parameter $\lambda_3^*$ controls any potential negative association between the gene counts. Specifically, when $\lambda_3^* = 0$, the counts are independent and when $\lambda_3^* < 0$ there is a negative association. [Footnote: We note that the original formulation of the OBPD model uses a slightly different parameterization in which the third parameter $0 \leq \lambda_3 \leq 1$ and where $\lambda_3 = 1$ indicates independence between counts. In order to ease interpretation of the model, we defined

$\lambda_3^* = \ln(\lambda_3)$ such that $-\infty < \lambda_3^* \leq 0$.] As $\lambda_3^*$ declines below 0, the negative association between counts becomes stronger and stronger.

The standard HBPD, which can detect positive associations between counts, also uses three parameters: $\lambda_1$ and $\lambda_2$ again represent the univariate Poisson rate parameters for KatE and KatG, respectively, when their counts are independent of one another. The third parameter $\lambda_3$ controls any potential positive association between the gene counts. Specifically, when $\lambda_3 = 0$, the counts are independent and when $\lambda_3 > 0$ there is a positive association. As $\lambda_3$ increases above 0, the positive association between counts becomes stronger and stronger.

To formally evaluate the hypothesis that there is strong selection for lineages to retain at least one copy of KatE or KatG, we formulated zero-modified extensions of both distributions. The zero-modified versions of these models are mixture models and introduce a fourth parameter $\delta_0$ that allows us to test whether the (0,0) counts were under- or over-represented relative to the rest of the data. Specifically, $P(0,0) = (1 - \delta_0) \cdot P_{BPD}(0,0)$. When $\delta_0 = 0$, the model reduces to the standard BPD. When $\delta_0 > 0$, fewer (0,0) organisms are observed than expected (zero-deflation), with $\delta_0$ directly interpretable as the fractional reduction in the (0,0) class probability. When $\delta_0 < 0$, more (0,0) organisms are observed than expected (zero-inflation). This extension decouples the (0,0) class, allowing us to directly test the hypothesis of additional selection for acquiring or retaining at least 1 Kat locus while still evaluating the relationship between KatE and KatG in the other gene count categories.

Because the standard OBPD and HBPD can be viewed as simpler, nested versions of our zero-modified mixture models, for either distribution we were able to evaluate all of the possible model combinations of shared and independent parameters within a classical, likelihood framework. The models were fitted using custom likelihood functions (https://github.com/mikegilchrist/Glasgo-Zinser) and the **bbmle** package (Bolker 2023) in R (R Core Team 2026). Using, in parallel, likelihood ratio tests (LRT) and corrected Akaike Information Criteria (AICc) we systematically compared different model formulations to determine which models are best supported by the data.

For our LRT analysis, we started with the most complex, 8 parameter model and then progressively evaluated simpler models with fewer parameters, asking whether or not we could fail to reject the simpler models with fewer unique parameters (**Fig. S10**). When the $\chi^2$-test statistic had a probability $> 0.05$ of occurring, the simpler model could not be rejected and became our new 'null' model. Confidence intervals were computed using profile likelihoods. For the $\delta_0$ parameter, whose MLE can be near the boundary of [0, 1), profile likelihood CIs were supplemented with delta-method CIs computed on the $\log(1 - \delta_0)$ scale and back-transformed to give proper asymmetric intervals.

A model's AICc explicitly takes its number of parameters into account and AICc values are only meaningful in the context of other AICc values for the same dataset. As a result, we calculated the *Δ*AICc for each model where, by convention the best fitting model has a value of *Δ*AICc $= 0$, and alternative models have values of *Δ*AICc $> 0$. The *Δ*AICc framework eschews formal statistical hypothesis testing and, instead, models with *Δ*AICc $\leq 2$ are considered to have similar support as the best fitting model.

Both model selection by AICc and the LRT identified the same best-supported model structure, though they favored slightly different model variants (see below). This minor discrepancy is expected given the different ways each framework penalizes model complexity, and the overall pattern of support is consistent across both approaches. Parameter estimates, confidence intervals, and likelihood ratio test statistics for the best-supported models are given in **Table S3**.

Under the zero-modified OBPD, the (0,0) count class was significantly under-represented relative to the standard OBPD prediction in both taxa ($\chi_2^2$ = 316.80, p < 1.6e-69): $\delta_0$ = 0.949 for *Vibrio* and $\delta_0$ = 0.472 for Proteobacteria (**Table S3**). This indicates that species lacking both enzymes are substantially rarer than expected given the other count data, suggesting additional selective pressure to maintain at least one catalase type. The zero-deflation is substantially stronger in *Vibrio* than in Proteobacteria.

Once we took the zero-modification into account, we found no evidence that KatE and KatG counts are positively or negatively associated under either model (Obrechkoff: $\lambda_3^*$ = -0.0087, $\chi_1^2$ = 0.25, p = 0.62; Holgate: $\lambda_3$ = 0.0000, $\chi_1^2$ = 0.00, p = 1.00). The best-supported OBPD model (**Table S3)** has KatE and KatG counts statistically independent ($\lambda_3^*$ = 0).

The KatE occurrence rate ($\lambda_1$) for Proteobacteria ($\lambda_1$ = 1.025) is substantially higher than for *Vibrio* ($\lambda_1$ = 0.506), while the KatG rate is similar across taxa ($\lambda_2$ = 0.617; **Table S3**). For *Vibrio*, KatE and KatG occur at comparable rates, whereas in Proteobacteria, KatE occurs at a substantially greater rate than KatG. A second model within $\Delta$AICc $\leq$ 2 additionally constrained *Vibrio* $\lambda_1$ = $\lambda_2$ (i.e., equal KatE and KatG rates), consistent with the similar KatE and KatG rates observed for *Vibrio*. The inability to distinguish between $\lambda_1$ and $\lambda_2$ for *Vibrio* likely reflects the much smaller sample size (n = 122) compared to Proteobacteria (n = 6,110). The final set of unique vs. shared parameter patterns are consistent with the visual trends in **Figures 2** and **S1**.

These results highlight the importance of biologically informed model fitting. If we had used the standard OBPD, which lacks the $\delta_0$ parameter for the (0,0) class, model fitting supports a significant negative association between KatE and KatG counts. By extending the model to account for additional selection against loss of all catalase activity, the mixture model reveals that the apparent negative association was driven entirely by the under-representation of the (0,0) class rather than a true covariance between the two enzyme types. Put another way, the standard OBPD conflates zero-deflation in the (0,0) class with negative covariance because a negative association is one way the model can ‘explain’ the reduced (0,0) counts. The mixture model separates these effects.

This analysis using a zero-modified extension of Obrechkoff bivariate Poisson mixture model illustrates how accounting for additional biological complexity not present in the standard BPD can fundamentally change one’s interpretation of the data. By recognizing the fundamentally different biological implications of going from zero catalases to one catalase vs. one catalase to two, we go from inferring a negative association between katE and katG copy numbers to independence of katE and katG copy numbers. Further, our model extension allows us to quantify the reduction in expected (0,0) count class, particularly among *Vibrio* species ($\delta_0 \approx 0.95$ for *Vibrio* vs. $\delta_0 \approx 0.47$ for Proteobacteria). Keeping our caveat about ignoring shared ancestry in our data analyses in mind and noting that the Poisson distribution also assumes independence between events, our results weakly support the interpretation that there is strong selection against loss of all catalase activity and a lack of evidence for selection favoring increased catalase copy numbers beyond just 1.

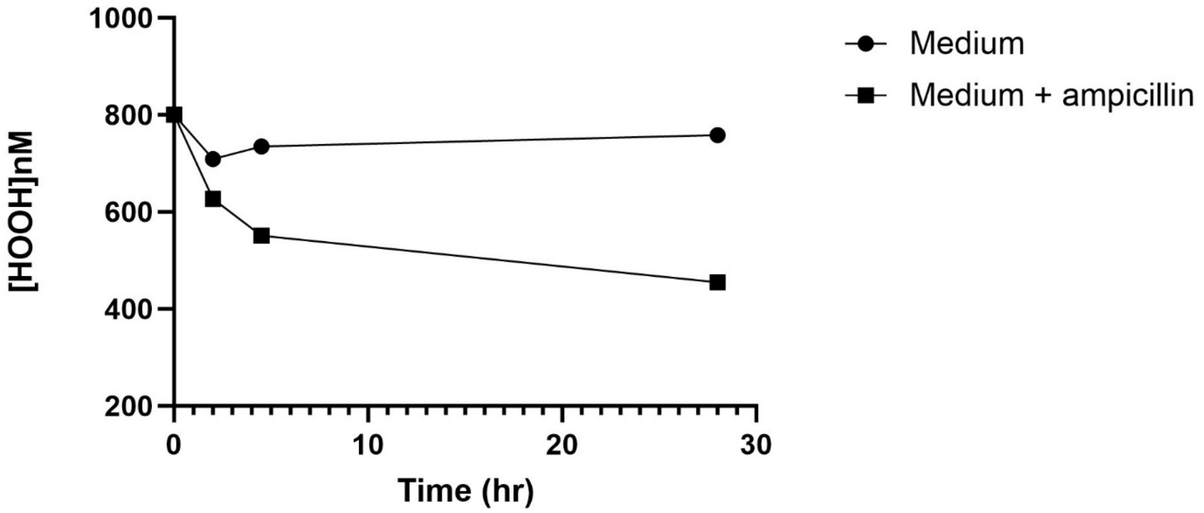


**Fig. S1.** Ampicillin-dependent decrease in hydrogen peroxide detection. (n=3; ± SD of the geometric mean).

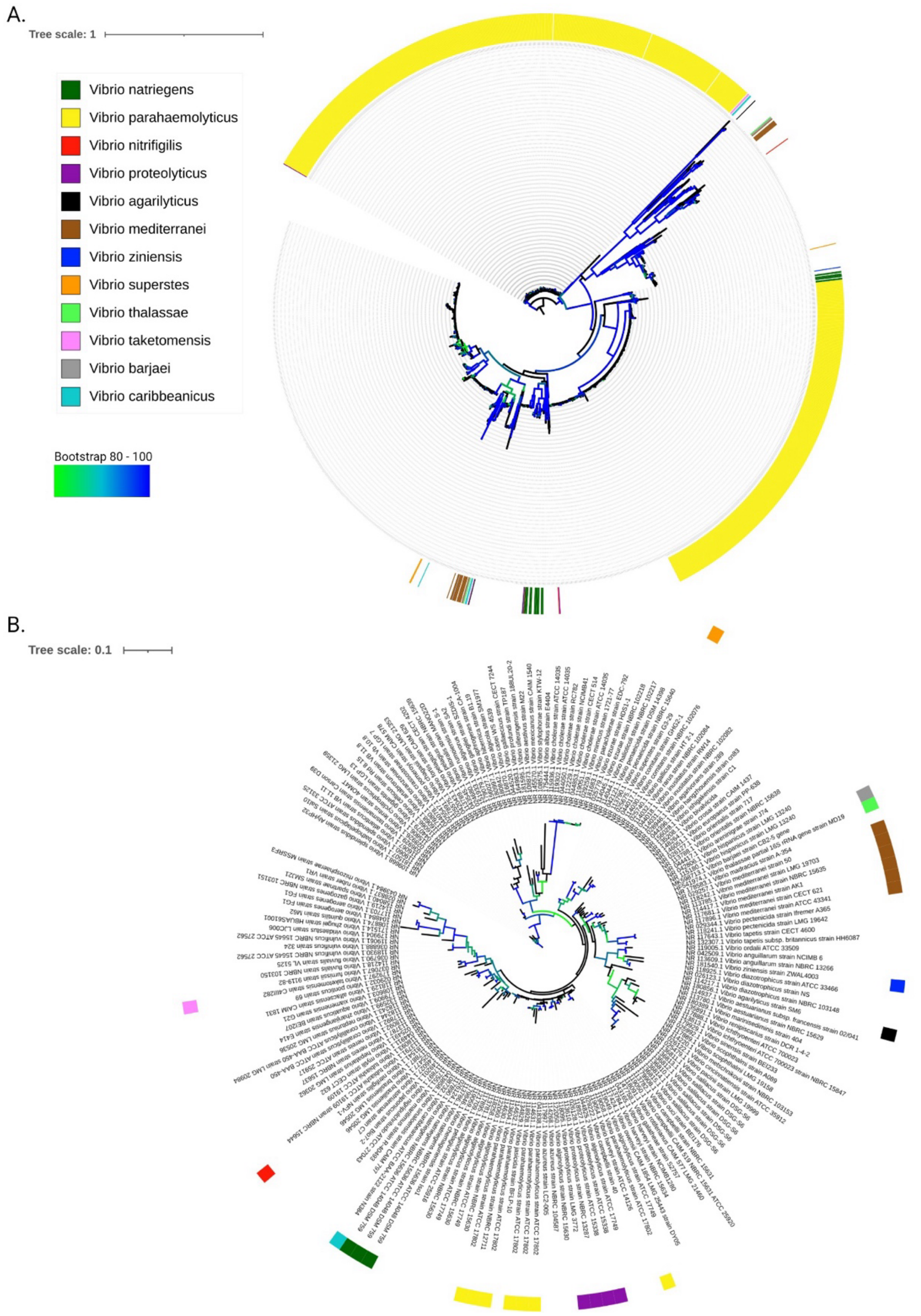


**Fig. S2.** (A) Uncollapsed *Vibrio spp.* KatG based phylogeny tree. (B) *Vibrio spp. 16S rRNA* based phylogeny tree. Species containing greater than one KatG (PF00141.26) copies are indicated by color strips. Branches with bootstrap values between 80 (green) to 100 (blue) are indicated.

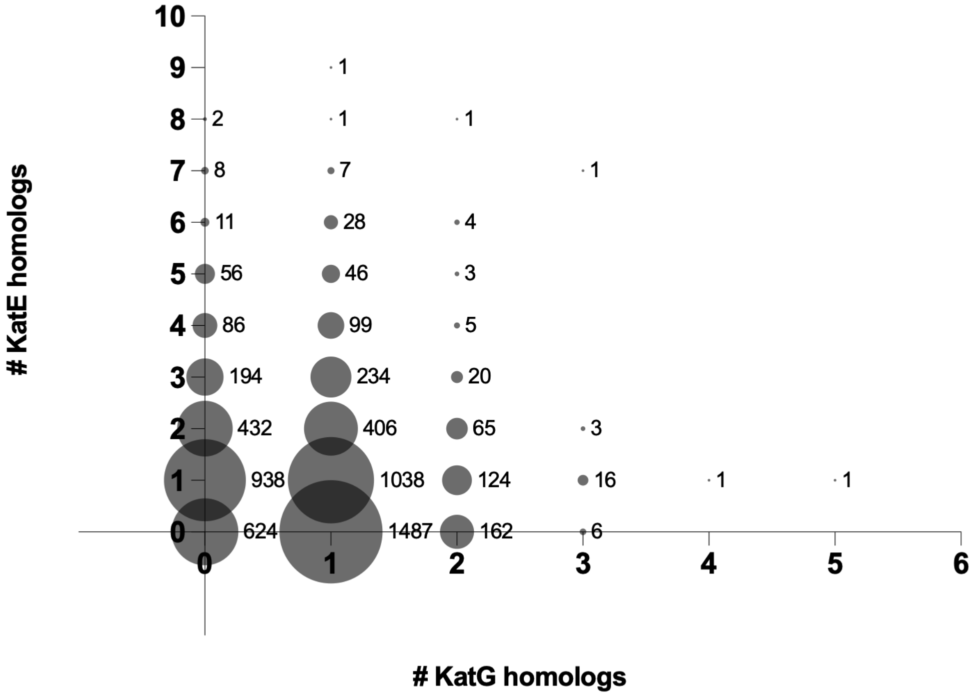


**Fig. S3.** Enumeration of KatE (PF00199.22) and KatG (PF00141.26) within Proteobacteria proteomes (Refseq proteomes accessed February 2022). ND = no data.

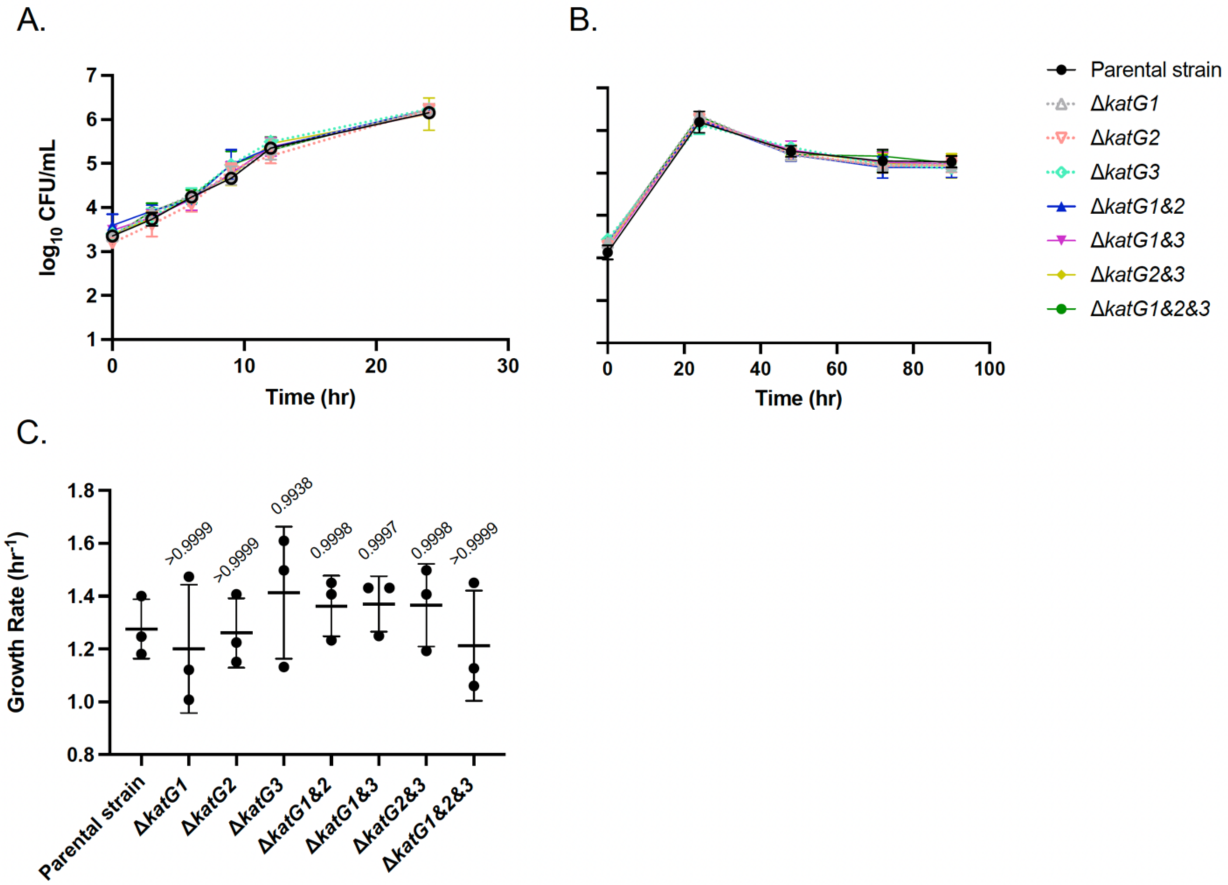


**Fig. S4.** *V. natriegens* parental and mutant strains growth curves in MHM + 24 µM acetate. (A and B) viability counts, (C) growth rates (n=3; ± SD of the geometric mean). Growth rates were calculated from the regression of cell numbers over three consecutive time points during exponential growth. P-values were calculated using a One-Way ANOVA multiple comparisons test with Dunnett's correction.

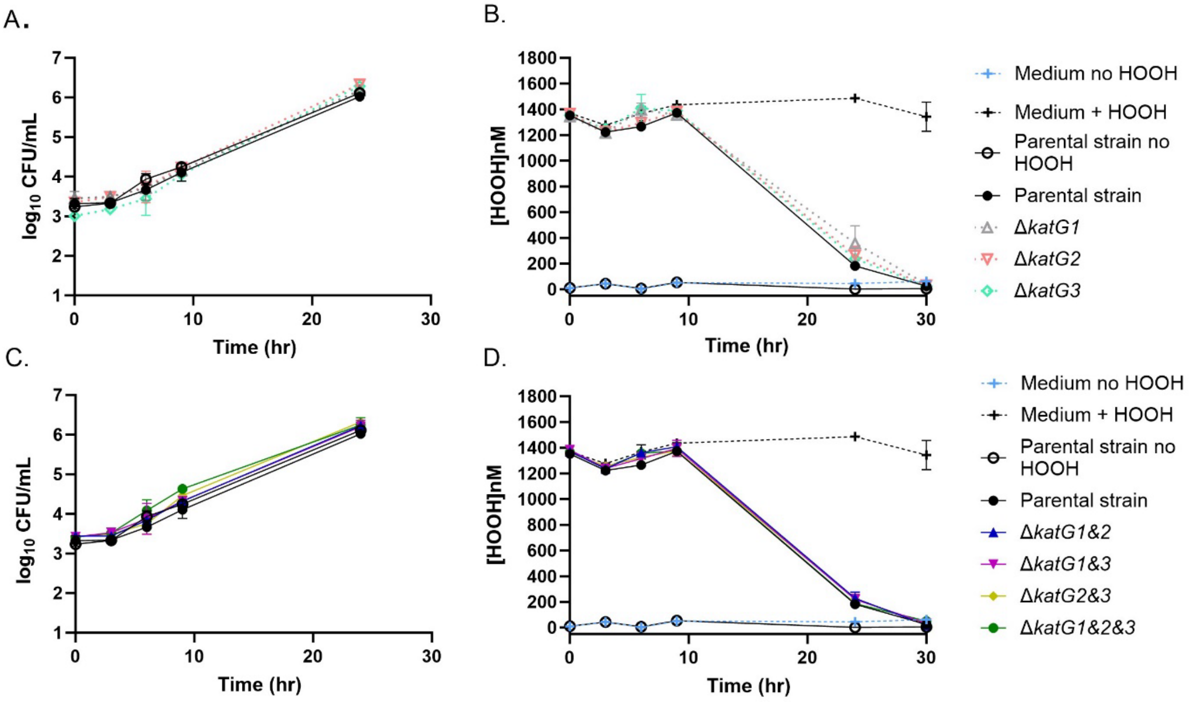


**Fig. S5.** (A and C) Cell viability and (B and D) HOOH degradation by *V. natriegens* parental and mutant strains in MHM + 24 µM acetate, (n=3; ± SD of the geometric mean). HOOH exposure at time 0.

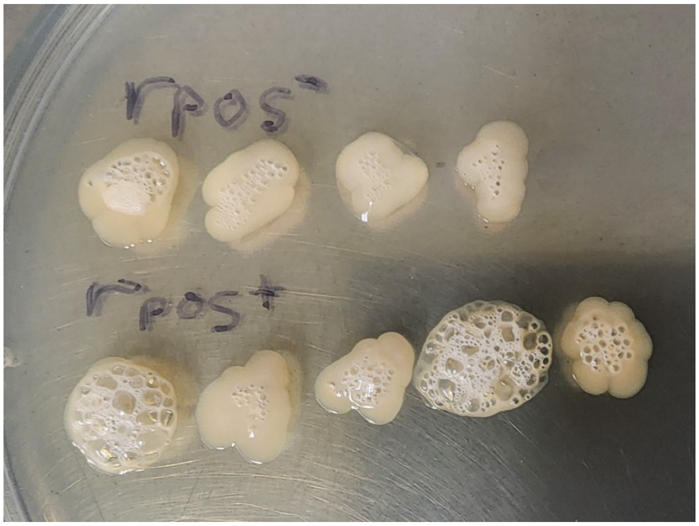


**Fig. S6.** Bubbling assay >5 minutes after HOOH exposure. Genotypes from left to right; *rpoS*$^{-}$ : 1) parental 2) Δ*katG1&2* 3) Δ*katG1&3* 4) Δ*katG2&3*. *rpoS*$^{+}$ : 1) parental *rpoS*E71 2) Δ*katG1&2 rpoS*E71 3) Δ*katG1&3 rpoS*E71 4) Δ*katG2&3 rpoS*E71 5) Δ*katG2&3 rpoS*E71N123.

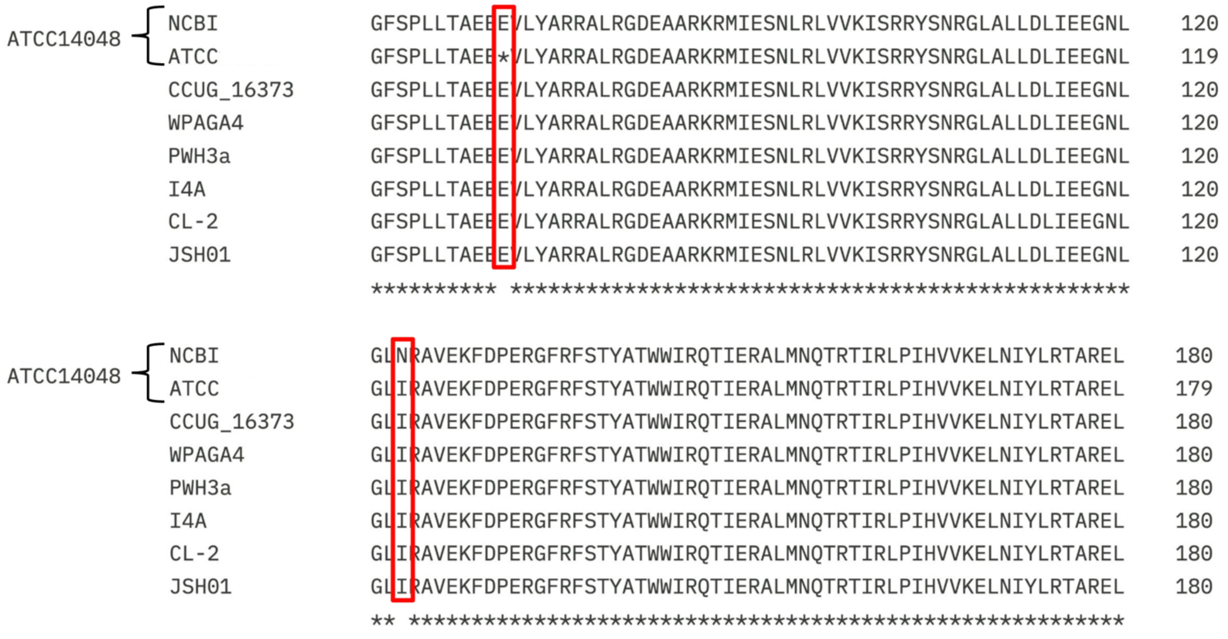


**Fig. S7.** Amino acid alignments of RpoS sequence segments from *Vibrio natriegens* ATCC 14048 and six geographically diverse isolates (CCUG 16371, WPAGA4, PWH3a, I4A, CL-2, and JSH01). Residues differing from the consensus sequence are highlighted in red boxes. Shown for the laboratory reference strain ATCC 14048 are sequences reported at either the NCBI or ATCC databases, revealing either an asparagine substitution at position 123 premature or a stop codon at amino acid 71, respectively. Notable, both deviations from consensus are absent from all six of the environmental isolates.

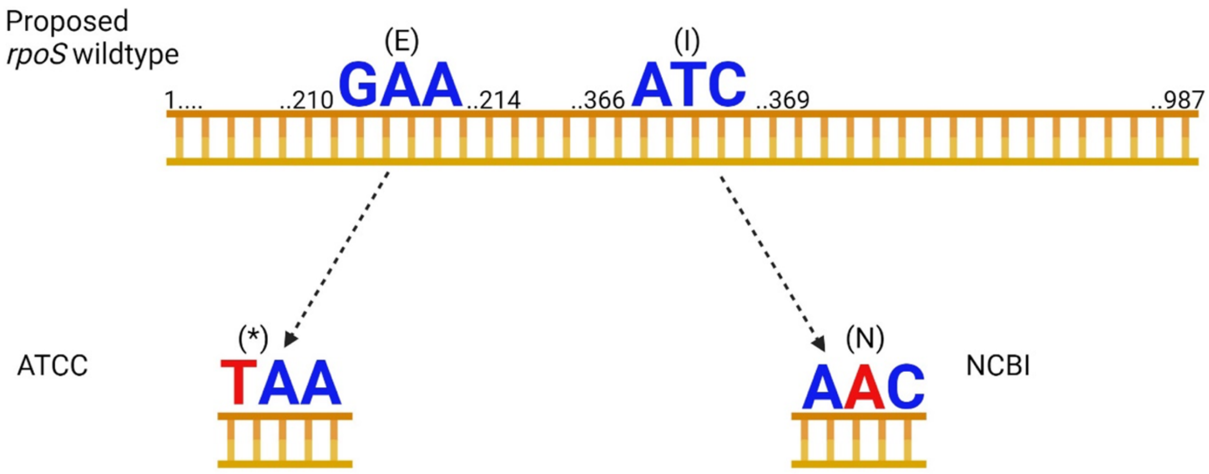


**Fig. S8.** Illustration of inferred ATCC14048 *rpoS* genotype history. (Top) Proposed "wildtype" *rpoS* ORF with nucleotide numbers given in black text. Codons found to have sequence variants are enlarged with colored text. Letters in red indicate the nucleotides proposed as mutations. Symbols inside brackets indicate the encoded amino acid (E, I, and N) or stop codon (*). (Bottom) Codon variants found in the *rpoS* sequence of indicated strain sequences. Created with BioRender.com.

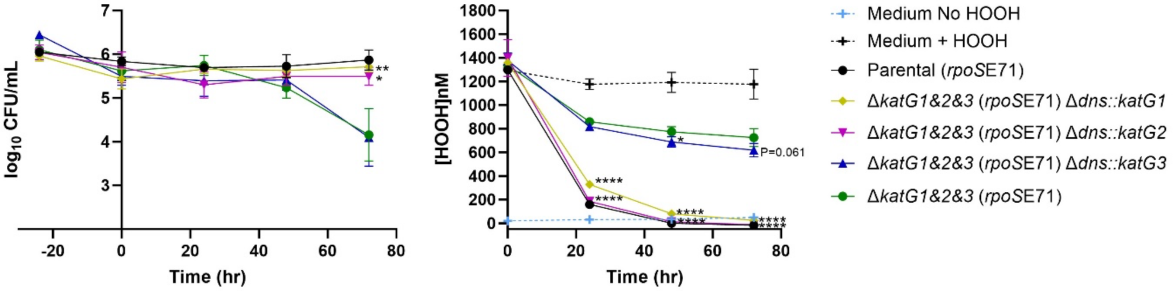


**Fig. S9.** Complementation of *katG* genes. (A) Cell viability and (B) HOOH degradation, (n=3; ± SD of the geometric mean). All genotypes have the E71I123 *rpoS* allele abbreviated as *rpoS*E71. HOOH exposure at time 0, after 24 hours of starvation in MHM + no carbon. P-values were calculated using a One-way ANOVA multiple comparisons test with Dunnett's correction, * $P \leq 0.05$, **** $P \leq 0.0001$.

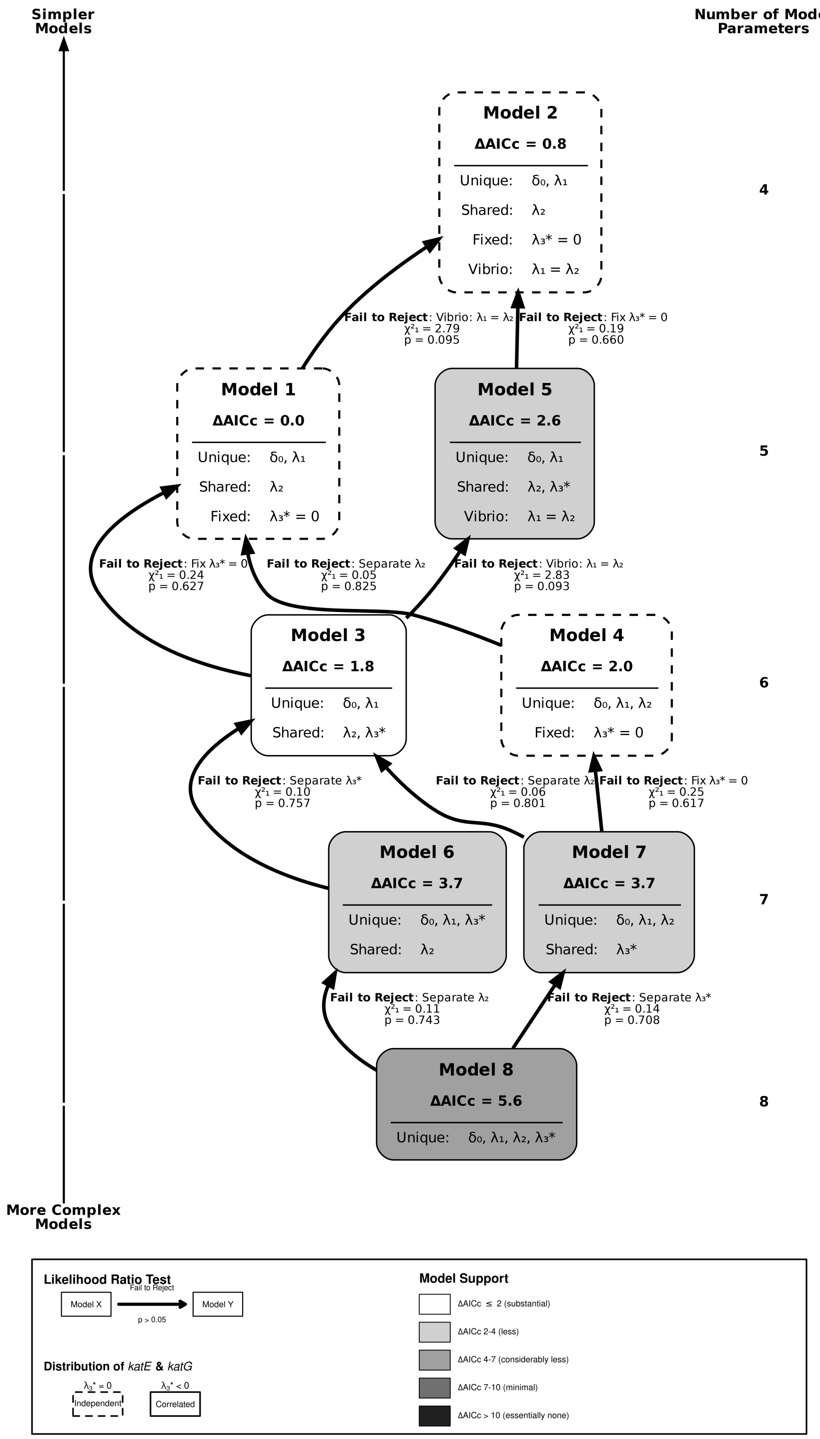


**Fig S10.** Summary of model selection pathway for arriving at the models best supported by the data (Models 1-4) for the Obrechkoff zero-modified bivariate Poisson mixture model, showing only constraints supported by likelihood ratio tests (LRT). Each node represents a model, labeled with its ΔAICc value and parameter structure (Unique = taxon-specific, Shared = common across taxa, Fixed = constrained to a boundary value). Arrows indicate nested model comparisons; solid arrows denote constraints that could not be rejected ($p > 0.05$), and dashed arrows denote rejected constraints. Models are arranged vertically by number of free parameters (simpler models at top, more complex at bottom). Node shading reflects ΔAICc: darker shading indicates poorer fit relative to the best model (ΔAICc = 0). The legend at bottom explains shading, arrow styles, and parameter abbreviations.

| Strain Name | Description | Source |
| --- | --- | --- |
| *Vibrio natriegens* ATCC 14048 pMMB*tfox* TND1964 (EZ260) | $Amp^R$ | Glasgo et al., 2024 |
| EZ262 | Chr.1 Δ1,520,008-1,520,031::$erm^R$, $Amp^R$ | Glasgo et al., 2024 |
| EZ264 | *ΔkatG1,* Chr.1 Δ1,520,008-1,520,031::$erm^R$, $Amp^R$ | This study |
| EZ265 | EZ260 *ΔkatG2,* Chr.1 Δ1,520,008-1,520,031::$erm^R$, $Amp^R$ | This study |
| EZ266 | *ΔkatG3,* Chr.1 Δ1,520,008-1,520,031::$erm^R$, $Amp^R$ | This study |
| EZ267 | *ΔkatG1, ΔkatG2,* Chr.1 Δ1,520,008-1,520,031::$erm^R$, $Amp^R$ | This study |
| EZ268 | *ΔkatG1, ΔkatG3,* Chr.1 Δ1,520,008-1,520,031::$erm^R$, $Amp^R$ | This study |
| EZ269 | *ΔkatG2, ΔkatG3,* Chr.1 Δ1,520,008-1,520,031::$erm^R$, $Amp^R$ | This study |
| EZ270 | *ΔkatG1, ΔkatG2, ΔkatG3,* Chr.1 Δ1,520,008-1,520,031::$erm^R$, $Amp^R$ | This study |
| EZ274 | *ΔoxyR,* Chr.1 Δ1,520,008-1,520,031::$erm^R$, $Amp^R$ | This study |
| EZ275 | *ΔkatG1, ΔkatG2, ΔkatG3, ΔoxyR* , Chr.1 Δ1,520,008-1,520,031::$erm^R$, $Amp^R$ | This study |
| EZ296 | *rpos*E71, Chr.1 Δ1,520,008-1,520,031::$erm^R$, $Amp^R$ | This study |
| EZ297 | *ΔkatG1, ΔkatG2, rpoS*E71, Chr.1 Δ1,520,008-1,520,031::$spec^R$, $Amp^R$ | This study |
| EZ298 | *ΔkatG1, ΔkatG3, rpoS*E71, Chr.1 Δ1,520,008-1,520,031::$spec^R$, $Amp^R$ | This study |
| EZ299 | *ΔkatG2, ΔkatG3, rpoS*E71, Chr.1 Δ1,520,008-1,520,031::$spec^R$, $Amp^R$ | This study |
| EZ300 | *ΔkatG2, ΔkatG3*, *rpoS*E71N123 , Chr.1 Δ1,520,008-1,520,031::$erm^R$, $Amp^R$ | This study |
| EZ308 | *ΔkatG1, ΔkatG2, ΔkatG3, rpoS*E71, Chr.1 Δ1,520,008-1,520,031::$spec^R$, $Amp^R$ | This study |
| EZ309 | *ΔkatG1, ΔkatG2, ΔkatG3, rpoS*E71N123, Chr.1 | This study |

| | | |
|---|---|---|
| | Δ1,520,008-1,520,031::$spec^R$, $Amp^R$ | |
| EZ310 | *ΔkatG1, ΔkatG2, ΔkatG3, rpoS*E71, Chr.1 Δ1,520,008-1,520,031::$erm^R$, $Amp^R$<br><br>Δ*dns*::*katG1* | This study |
| EZ311 | *ΔkatG1, ΔkatG2, ΔkatG3, rpoS*E71, Chr.1 Δ1,520,008-1,520,031::$erm^R$, $Amp^R$<br><br>Δ*dns*::*katG2* | This study |
| EZ312 | *ΔkatG1, ΔkatG2, ΔkatG3, rpoS*E71, Chr.1 Δ1,520,008-1,520,031::$erm^R$, $Amp^R$<br><br>Δ*dns*::*katG3* | This study |

**Table S1.** Strains used in this study.

| Primer name | 5' -> 3' sequence | Description |
|---|---|---|
| katG1F1 | CCAATTGGTGGAGAGAGTGC | *ΔkatG1* Upstream forward primer |
| katG1R1 | GCTAATTCAGTTTAAGCGGCCATCTCCTTTGTTAGGGGCACAG | *ΔkatG1* Upstream reverse primer |
| katG1F2 | ATGGCCGCTTAAACTGAATTAGCCAAACGAGTTCTCTTAATTAACTGTC | *ΔkatG1* downstream forward primer |
| katG1R2 | CTCGATGCAGGCGCTATT | *ΔkatG1* downstream reverse primer |
| 133308F | TCTGTCGAGCTGATGCAAGT | *ΔkatG1* forward scar primer |
| 136180R | AGAACCAATCATCGGGTTTG | *ΔkatG1* reverse scar primer |
| katG2F1 | AGCTTCATCTTTTGCCATGC | *ΔkatG2* Upstream forward primer |
| katG2R1 | GCTAATTCAGTTTAAGCGGCCATGCTGTTGTTCATAGGGTGTCC | *ΔkatG2* Upstream reverse primer |
| katG2F2 | ATGGCCGCTTAAACTGAATTAGCGAAATGCACTTTGGTGCGTA | *ΔkatG2* downstream forward primer |
| katG2R2 | CGCAGGACAACTGGAAAATA | *ΔkatG2* downstream reverse primer |
| 447588F | GCACACGTTATCACCTCTCCT | *ΔkatG2* forward scar primer |
| 450432R | AAAGAGAGCGCTGACCTGAG | *ΔkatG2* reverse scar primer |
| katG3F1 | ATAGCCACCATCGGTTGAGA | *ΔkatG3* Upstream |

| | | |
|---|---|---|
| | | forward primer |
| katG3R1 | GCTAATTCAGTTTAAGCGGCCATCTGGCTTTTTGCGTATCCAT | *ΔkatG3* Upstream reverse primer |
| katG3F2 | ATGGCCGCTTAAACTGAATTAGCTCGCTAAGCTAGCATAAGCTCT | *ΔkatG3* downstream forward primer |
| katG3R2 | GGATTCTGACTGGAGCAAGC | *ΔkatG3* downstream reverse primer |
| 1119752F | GCCAGACATCCTAATGCCTTT | *ΔkatG3* forward scar primer |
| 1122735R | GTTCCGGAGATATCGGACAA | *ΔkatG3* reverse scar primer |
| endAF1 | CTAACATGGCTAAGCACCTG | *Δdns* upstream forward primer |
| endAR1 | ACACAATCGCTCAAGACGTGACTGAGGATTAGGAAAGCTGGA | *Δdns* upstream reverse primer |
| endAF2 | ATGGCCGCTTAAACTGAATTAGCCCTCACCAATCGCGACAATC | *Δdns* downstream forward primer |
| endAR2 | TAAGGTGTCTCAAATCTCAATCTAGG | *Δdns* downstream reverse primer |
| KatG1compF | CACGTCTTGAGCGATTGTGTAAATCATCAGCAATCTGGTGT | *katG1* complement forward primer |
| KatG1compR | GCTAATTCAGTTTAAGCGGCCATGAGTCCGCCTGAATTTATCG | *katG1* complement reverse primer |
| KatG2compFH1 | CACGTCTTGAGCGATTGTGTCCCGAGCTCCAACAATAGAT | *katG2* complement |

| | | |
|---|---|---|
| | | forward primer |
| KatG2compRH3 | GCTAATTCAGTTTAAGCGGCCATGAGTCCGCCTGAATTTATCG | *katG2* complement reverse primer |
| KatG3compFH1 | CACGTCTTGAGCGATTGTGAATGTCTGGCATGGATACGC | *katG3* complement forward primer |
| KatG3compRH3 | GCTAATTCAGTTTAAGCGGCCATGCCCATCTTCCTATTGATTCTG | *katG3* complement reverse primer |
| 2728836F | TTCCCTATTCCCAGCCTGAC | *Δdns::katG s*car forward primer |
| 2730429R | CTACGCGCTCAGAGATGTCT | *Δdns::katG s*car reverse primer |
| rpoSF1 | TCGTTTCTGCGCGTTTAGAG | *rpoS* upstream forward primer |
| rpoSE71R1 | AAGCACTTCTTCTTCAGCAGT | *rpoS* upstream reverse primer |
| rpoSE71F2 | ACTGCTGAAGAAGAAGTGCTT | *rpoS* middle piece forward primer |
| rpoSN123R2 | TCTCAACTGCACGGTTCAAG | *rpoS* middle piece reverse primer |
| rpoSN123F3 | CTTGAACCGTGCAGTTGAGA | *rpoS* downstream forward primer |
| rpoSR3 | TTCACGTATCCTAAGCCGCT | *rpoS downstream* reverse primer |
| RpoSF | TGTCATGTCTTGCTAACTCGC | *rpoS* ORF forward primer |

| | | |
|---|---|---|
| RpoSR | TTGGATACCCTGCTTAAAACGC | *rpoS* ORF reverse primer |

**Table S2.** Primer sequences used in this study. Underlined nucleotides indicate homologous regions added for SOE PCR.

| Model | npar | ΔAICc | Taxon | $\delta_0$ | $\lambda_1$ | $\lambda_2$ | $\lambda_3$* |
|---|---|---|---|---|---|---|---|
| 1 | 5 | 0.0 | Vibrio | 0.95 (0.80 – 0.99) | 0.51 (0.38 – 0.63) | **0.62 (0.58 – 0.65)** | **0†** |
| | | | Proteo | 0.47 (0.42 – 0.52) | 1.02 (0.98 – 1.07) | **0.62 (0.58 – 0.65)** | **0†** |
| 2 | 4 | 0.8 | Vibrio | 0.94 (0.78 – 0.99) | 0.62‡ (0.58 – 0.65) | **0.62 (0.58 – 0.65)** | **0†** |
| | | | Proteo | 0.47 (0.43 – 0.52) | 1.02 (0.98 – 1.07) | **0.62 (0.58 – 0.65)** | **0†** |
| 3 | 6 | 1.8 | Vibrio | 0.95 (0.80 – 0.99) | 0.51 (0.39 – 0.63) | **0.62 (0.59 – 0.66)** | **-0.01 (-0.04 – 0.03)** |
| | | | Proteo | 0.47 (0.42 – 0.52) | 1.03 (0.99 – 1.07) | **0.62 (0.59 – 0.66)** | **-0.01 (-0.04 – 0.03)** |
| 4 | 6 | 1.9 | Vibrio | 0.95 (0.80 – 0.99) | 0.50 (0.37 – 0.63) | 0.60 (0.45 – 0.75) | **0†** |
| | | | Proteo | 0.47 (0.42 – 0.52) | 1.02 (0.98 – 1.07) | 0.62 (0.58 – 0.65) | **0†** |
| 5 | 5 | 2.6 | Vibrio | 0.94 (0.78 – 0.99) | 0.62‡ (0.59 – 0.66) | **0.62 (0.59 – 0.66)** | **-0.01 (-0.04 – 0.03)** |
| | | | Proteo | 0.47 (0.42 – 0.52) | 1.03 (0.99 – 1.07) | **0.62 (0.59 – 0.66)** | **-0.01 (-0.04 – 0.03)** |
| 6 | 7 | 3.7 | Vibrio | 0.95 (0.80 – 0.99) | 0.51 (0.37 – 0.65) | **0.63 (0.59 – 0.66)** | 0.00 (-0.22 – 0.22) |
| | | | Proteo | 0.47 (0.42 – 0.52) | 1.03 (0.99 – 1.08) | **0.63 (0.59 – 0.66)** | -0.01 (-0.04 – 0.02) |
| 7 | 7 | 3.7 | Vibrio | 0.95 (0.80 – 0.99) | 0.50 (0.37 – 0.64) | 0.60 (0.46 – 0.75) | **-0.01 (-0.04 – 0.03)** |
| | | | Proteo | 0.47 (0.42 – 0.52) | 1.03 (0.99 – 1.07) | 0.62 (0.59 – 0.66) | **-0.01 (-0.04 – 0.03)** |
| 8 | 8 | 5.6 | Vibrio | 0.95 (0.80 – 0.99) | 0.50 (0.30 – 0.70) | 0.60 (0.38 – 0.82) | 0.00 (-0.33 – 0.33) |
| | | | Proteo | 0.47 (0.42 – 0.52) | 1.03 (0.99 – 1.08) | 0.63 (0.59 – 0.66) | -0.01 (-0.05 – 0.02) |

**Table S3**: Parameter estimates (95% CI) for top 8 models by ΔAICc. Each model has two rows (Vibrio, Proteobacteria). npar = number of model parameters. Bold = shared across taxa. † = fixed at boundary value. ‡ = constrained equal to $\lambda_2$.

| Model | npar | Unique | Shared | Fixed | Equality | log L | ΔAICc |
|---|---|---|---|---|---|---|---|
| **1** | **5** | **$\delta_0$, $\lambda_1$** | **$\lambda_2$** | **$\lambda_3$* = 0** | **—** | **-11910.47** | **0.00** |
| **2** | **4** | **$\delta_0$, $\lambda_1$** | **$\lambda_2$** | **$\lambda_3$* = 0** | **Vibrio: $\lambda_1 = \lambda_2$** | **-11911.86** | **0.78** |
| **3** | **6** | **$\delta_0$, $\lambda_1$** | **$\lambda_2$, $\lambda_3$*** | **—** | **—** | **-11910.35** | **1.77** |
| **4** | **6** | **$\delta_0$, $\lambda_1$, $\lambda_2$** | **—** | **$\lambda_3$* = 0** | **—** | **-11910.44** | **1.95** |
| 5 | 5 | $\delta_0$, $\lambda_1$ | $\lambda_2$, $\lambda_3$* | — | Vibrio: $\lambda_1 = \lambda_2$ | -11911.76 | 2.59 |
| 6 | 7 | $\delta_0$, $\lambda_1$, $\lambda_3$* | $\lambda_2$ | — | — | -11910.30 | 3.68 |
| 7 | 7 | $\delta_0$, $\lambda_1$, $\lambda_2$ | $\lambda_3$* | — | — | -11910.32 | 3.71 |
| 8 | 8 | $\delta_0$, $\lambda_1$, $\lambda_2$, $\lambda_3$* | — | — | — | -11910.25 | 5.57 |
| 9 | 7 | $\delta_0$, $\lambda_2$, $\lambda_3$* | $\lambda_1$ | — | — | -11921.02 | 25.12 |
| 10 | 6 | $\lambda_2$, $\lambda_3$* | $\delta_0$, $\lambda_1$ | — | — | -11926.82 | 34.72 |
| 11 | 6 | $\delta_0$, $\lambda_3$* | $\lambda_1$, $\lambda_2$ | — | — | -11927.26 | 35.58 |
| 12 | 5 | $\delta_0$, $\lambda_2$ | $\lambda_1$ | $\lambda_3$* = 0 | — | -11928.67 | 36.40 |
| 13 | 4 | $\delta_0$ | $\lambda_1$, $\lambda_2$ | $\lambda_3$* = 0 | — | -11930.20 | 37.47 |
| 14 | 6 | $\delta_0$, $\lambda_2$ | $\lambda_1$, $\lambda_3$* | — | — | -11928.49 | 38.04 |
| 15 | 5 | $\delta_0$ | $\lambda_1$, $\lambda_2$, $\lambda_3$* | — | — | -11930.02 | 39.10 |
| 16 | 4 | $\lambda_2$ | $\delta_0$, $\lambda_1$ | $\lambda_3$* = 0 | — | -11934.22 | 45.50 |
| 17 | 5 | $\lambda_2$ | $\delta_0$, $\lambda_1$, $\lambda_3$* | — | — | -11934.04 | 47.14 |
| 18 | 3 | -- | $\delta_0$, $\lambda_1$, $\lambda_2$ | $\lambda_3$* = 0 | — | -11937.39 | 49.85 |
| 19 | 5 | $\lambda_3$* | $\delta_0$, $\lambda_1$, $\lambda_2$ | — | — | -11935.51 | 50.09 |
| 20 | 4 | -- | $\delta_0$, $\lambda_1$, $\lambda_2$, $\lambda_3$* | — | — | -11937.21 | 51.48 |

**Table S4**: Model comparison for all 20 Obrechkoff zero-modified bivariate Poisson mixture models. Unique = taxon-specific parameters; Shared = common across taxa; Fixed = constrained to boundary value; Equality = within-taxon parameter constraints. npar = number of model parameters. Bold rows indicate models with ΔAICc ≤ 2 (substantial support).

**Supplemental Video 1: Bubbling assay of HOOH-exposed strains in real time.**
Genotypes from left to right; *rpoS*$^{-}$ : 1) parental 2) Δ*katG1&2* 3) Δ*katG1&3* 4) Δ*katG2&3*. *rpoS*$^{+}$ : 1) parental *rpoS*E71 2) Δ*katG1&2 rpoS*E71 3) Δ*katG1&3 rpoS*E71 4) Δ*katG2&3 rpoS*E71 5) Δ*katG2&3 rpoS*E71N123.
